# A compressed code for memory discrimination

**DOI:** 10.1101/2025.10.12.681901

**Authors:** Dale Zhou, Sharon M. Noh, Nora C. Harhen, Nidhi V. Banavar, Catherine A. Hartley, C. Brock Kirwan, Michael A. Yassa, Aaron M. Bornstein

## Abstract

The ability to discriminate similar visual stimuli has been used as an important index of memory function. This ability is widely thought to be supported by expanding the dimensionality of relevant neural codes, such that neural representations for the similar stimuli are maximally distinct, or “separated.” An alternative hypothesis is that discrimination is supported by lossy compression of visual inputs, efficiently coding sensory information by discarding seemingly irrelevant details. A benefit of compression, relative to expansion, is that it allows the individual to efficiently retain fewer essential dimensions underlying stimulus variation—a process linked to higher-order visual processing—without hindering discrimination. Under the compression hypothesis, memory discrimination is facilitated when more redundant information from similar stimuli can be discarded to emphasize distinct information. We test the compression versus expansion hypotheses by predicting performance on a memory discrimination task (combined *n* = 723). First, we train neural networks to compress perceptual and semantic factors of stimuli, and measure the lossiness of those representations using the mathematical framework underlying compression. Consistent with the compression hypothesis, we find that greater lossiness predicts the ease and performance of lure discrimination. We empirically confirm these predictions across image sets, four behavioral datasets, and alternative metrics of lossiness. Using representational similarity analysis and two task fMRI datasets (combined *n* = 91), we identify compressed representations in the higher-order visual stream inferotemporal cortex (IT) as well as hippocampal subregion dentate gyrus/CA3. Finally, we observe retrieval-related compression in biologically realistic neural network models of cortico-hippocampal pattern separation. These results suggest lossy compression may support memory discrimination.

## 1 Introduction

Many behaviors, from value-based decisions *(1–4)* and associative learning *(5)* to perceptual inference and memory *(6–10)*, require recognizing whether the perception of a current situation is familiar or novel. This process is challenging because memory is constructive. From a percept with partial information, memory integrates prior experiences to fill in gaps but, in doing so, introduces distortions that can create incorrect impressions of prior experience *(11, 12)*. Discriminating whether the current situation is novel compared to remembered experiences depends on pattern separation *(13–15)*. Pattern separation occurs when brain regions, such as the hippocampus, transform similar inputs that would produce similar activity patterns into output patterns that are more distinct **(Figure 1A)** *(16–26)*. Distinctness is often measured by the degree of linear independence where maximally distinct patterns are orthogonal.

**Figure 1.**
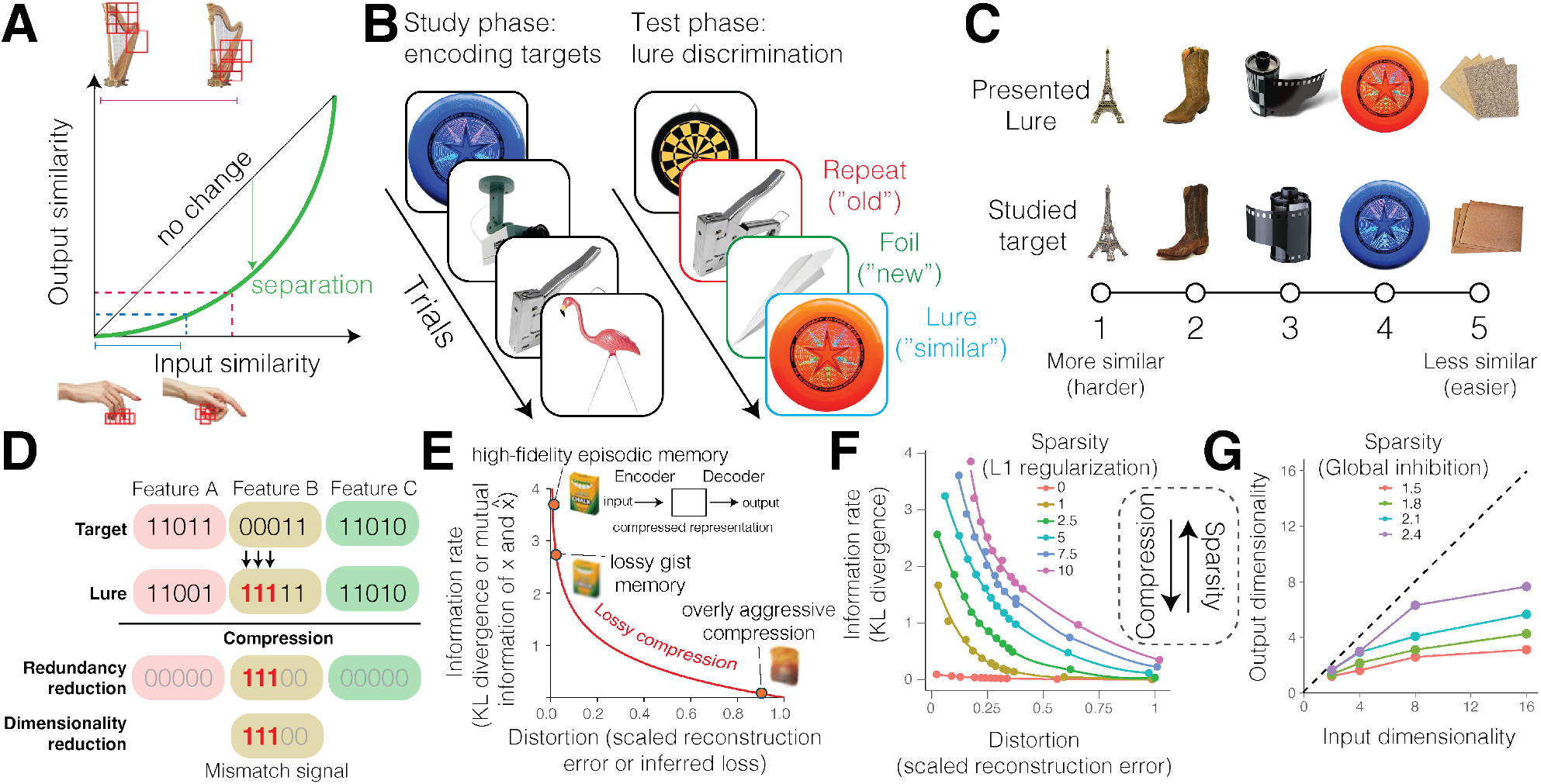
Task schematic and lossy compression function. **(A)** This canonical input-output function is the computational modeling target of pattern separation, encoding similar inputs into more dissimilar representations. Distinguishing between extremely similar harps may require encoding of more information (red-dashed lines) than distinguishing between more dissimilar hand gestures (blue-dashed lines). Red bounding boxes indicate patches of maximal importance for discrimination by the AlexNet neural network *(68)*. **(B)** Task stimuli during the study and test phases with corresponding trial type and correct response. **(C)** The ease of memory discrimination between target and lure images was based on prior behavioral performance *(69)*. **(D)** Hypothesized lossy compression computation supporting pattern separation. The stimulus and its lure are abstracted as one-hot representations for categorically structured stimuli. The lure is similar in all but the perceptual Feature B (difference highlighted in red). Compression retains the unique information in Feature B while discarding redundant information or feature dimensions. **(E)** Information theory provides insights into what statistics of the inputs can be discarded as redundant according to a rate-distortion function, the mathematical basis of compression. A high-fidelity memory might be associated with the semantic representation like: “a new yellow and green box of crayon chalk in white”. A moderately compressed memory loses some detail: “a new yellow and green box of crayons.” Finally, a very aggressively compressed memory loses essential details, resulting in false memory: “a used, crumpled yellow and red bag.” **(F)** We used computational model of compression, where *β*-variational autoencoders learn to reconstruct each task image using compressed probabilistic representations *(53, 63, 64, 70)*. The Kullback–Leibler (KL) divergence indicates the information rate allocated to encode images beyond a Gaussian prior, whereas the distortion is the reconstruction error. These information theory metrics discriminate between sparsity (by inhibiting the magnitude of encoding units through L1 regularization) and compression (i.e. allocating a reduced information rate). Increasing sparsity worsens compression. Data points are averages across images per *β*. **(G)** We used a biologically realistic neural network model of compression in a cortical layer of neurons operating under purely Hebbian plasticity to process inputs *(22, 71)*. Similarly to (F), dimensionality metrics distinguish between sparsity and compression. Increasing sparsity by raising the global inhibition mitigates dimensionality reduction. Points that fall further below the the dashed line (a reference line indicating the dimensionality of the output is the same as the input) indicate more compression.

The longstanding Marr-Albus hypothesis suggests two neural pathways to orthogonalize representations: expand the encoding ensemble and sparsen encoding activity *(27, 28)*. Both expanding the number of encoding units (neurons) using divergent projections from a small to a large population (a ratio of about 1:5 in the hippocampus) and inhibiting the population activity to have few active neurons within a relevant timespan (around 5% of neurons active) can decorrelate the statistical structure of inputs to separate across distinct ensembles of neurons *(5, 17, 19, 27–34)*. However, many cortical regions implicated in pattern separation lack the hippocampus’ distinctive expansion and sparsity *(13, 15, 23, 35–41)*.

A less explored hypothesis for pattern separation is lossy compression, which extends Barlow-Attneave’s redundancy reduction principle by discarding redundant information to produce efficient representations while tolerating some error *(42–45)*. Memory discrimination is supported by reducing redundant overlap and keeping distinct details between inputs *(13, 30, 46)*, but it is not understood how this redundancy reduction occurs. Compressed coding is one candidate. In contrast to the Marr-Albus expansion, compression suggests roles for reducing redundant information in an encoding ensemble by creating a physical or information bottleneck that encourages more compact representations. Such representations can become orthogonalized by retaining the most distinctive information. Taken together, the Marr-Albus expansion and Barlow-Attneave compression hypotheses make theoretically distinguishable predictions for how pattern separation could be multiply realizable across the brain.

Here, we test if lossy compression can explain memory discrimination in a behavioral task designed to tax pattern separation, the Mnemonic Similarity Task *(26)*. In a study phase, participants view a sequence of images, each showing an everyday object such as a pineapple, and judge whether the object belongs indoors or outdoors. This judgment serves as a cover task, so participants encode the “target” images incidentally from a single exposure to each. In a test phase, participants view individual images and classify each as a previously studied “repeat,” a similar but distinct “lure,” or a novel “foil,” which measures their ability to discriminate between similar memories **(Figure 1B)**. Memory discrimination, measured by the proportion of correctly classified lures, is harder when the lure image is more similar to the target image (**Figure 1C**). This ability to discriminate is thought to assess detail knowledge or specific recollection *(30, 47, 48)*.

We analyzed performance on 1,152 target-lure pairs across four previously published behavioral datasets (combined *n* = 723; see **Materials and Methods**) *(9, 49–51)* as well as two previously published fMRI measurements of brain activity during the task (combined *n* = 91) *(37, 38)*. Because visual information reaches the hippocampus after processing by visual and semantic cortical pathways *(15, 52)*, we operationalized various perceptual and/or semantic features that may ultimately become compressed. To do so, we process images through neural networks trained for pixel reconstruction, image classification, or image-to-text conversion *(53–57)*. Deep-layer representations in neural networks are more “semantic”-like because supervised learning across diverse object categories enables them to capture nameable properties accumulated across experiences *(58)*. In contrast, shallow-layer representations are more “perceptual”-like, primarily capturing lower-level visual features learned from far less and unlabeled data (see **Materials and Methods** for caveats to this operationalization). Such representations have been predictive of the neural activity of temporal and visual cortices *(59–61)*. Compression is operationalized by several convergent approaches. In behavioral data, we use an information-theoretic algorithm that we modify to estimate lossiness from a cosine similarity metric of orthogonalization between item pairs *(43)* and auto-associative networks that measure lossiness by individual item reconstruction errors *(53, 62–64)*. In neural data, we use representational similarity analysis (RSA) to test whether stimulus representations exhibit the hypothesized compression. We compress the neural network representations which were most predictive in the behavioral d atasets by training them on a self-supervised learning instantiation of Barlow-Attneave’s redundancy reduction objective *(65)* and manipulate the severity of an information bottleneck *(66)* to test different degrees of compression in neural representations. Finally, we investigate signatures of compression in a biologically realistic neural network model of cortico-hippocampal pattern separation function *(13, 67)*.

We hypothesize that memory discrimination is more difficult and worsened when more bits of information are needed to preserve high-fidelity information about subtle differences in detail between target and lure stimulus pairs **(Fig 1D)**. Conversely, memory discrimination is easier and better when lossy compression can discard more bits for greater efficiency. In contrast to compressing target-lure stimulus pairs, compressing each stimulus individually before knowing whether its information is unique or redundant relative to an unseen lure should produce systematic lure-discrimination errors. We hypothesize that aggressive compression of individual stimuli, resulting in more bits discarded per unit of lossiness, increases false alarms (**Figure 1E)**. We find evidence in support of the lossy compression hypothesis, replicated across image datasets, image features, compression methods, behavioral datasets, and neural datasets. Together, these findings suggest complementary computational roles for cortex and hippocampus in memory discrimination, with compressed cortical representations and hippocampal representations balancing sparse expansion with retrieval-related compression.

## 2 Results

### Theoretical distinctions between compression and expansion hypotheses

We hypothesize a novel computational pathway by which representations can become more dissimilar according to Barlow-Attneave’s efficient coding hypothesis of redundancy reduction. To highlight differences with the Marr-Albus model, we summarize how the two processes increase the dissimilarity of inputs, which is the shared modeling target for any theory of pattern separation *(72)*.

Theories of pattern separation must explain how the similarity between inputs *X*_*target*_ and *X*_*lure*_ decreases according to an arbitrary similarity metric, *S*(*X*_*target*_, *X*_*lure*_), such as cosine similarity *(72)*. Pattern separation occurs when *S*(*X*_*target*_, *X*_*lure*_) decreases across neural regions or neuronal populations *A, B*, and *C*, such that *X*_*target*_ and *X*_*lure*_ are progressively decorrelated:

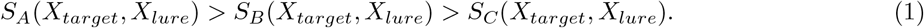

Expansion model (Marr-Albus sparse coding):

1. Dissimilarity arises from representations becoming sparser through inhibitory signaling and many efferent projections to separate neural units.
2. Separate neural units independently encode (orthogonalize) the lure and its associate, preserving their information in a greater number of dimensions.

Compression model (Barlow-Attneave efficient coding):

1. Dissimilarity arises from discarding redundant bits and dimensions of information between the lure and its associate, which results in selective distortion of the original inputs.
2. Selective distortion is useful for discrimination when it preserves unique and discards redundant information or dimensions.

The computational kinship between pattern separation and lossy compression can be made more apparent mathematically, distinguishing this account from established hypotheses such as sparsity. Although *S* is commonly conceived as a linear correlation like cosine similarity, *S* can also be defined as the mutual information *I* to capture both linear and nonlinear dependencies between inputs. Under this definition of *S*,

Equation 1 directly parallels the data processing inequality *(73)*:

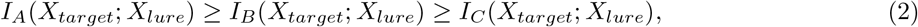

stating that physical processing from *A* → *B* → *C* cannot create new information about the original source. The inequality points to a trade-off where information is either retained at some cost or lost for more lightweight but error-prone transmission, a core computational problem of memory. Memory needs to reconstruct arbitrary traces yet cannot preserve all of the information and structure of inputs *(46)*.

How many bits of information should be allocated to more precise high-fidelity memory versus saved for more approximate gist memory *(63, 74)*? Lossy compression provides a framework to determine the optimal solution to this trade-off. Optimally, lossy compression is the joint minimization of (1) the *rate* of information *R* needed to encode an input *X*_*target*_ as a compressed representation *X*_*lure*_ and (2) the amount of *distortion D* caused by information lost from compressing *X*_*target*_ into *X*_*lure*_ *(75)*:

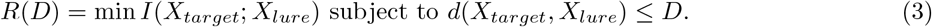

Rate-distortion theory shows how lossy compression forces approximations such that *D >* 0. We propose that reducing *S*(*X*_*target*_; *X*_*lure*_) for pattern separation involves minimizing existing redundancy, *I*(*X*_*target*_; *X*_*lure*_), which is marked by detectable increases in selective distortion *D*_*A*_(*X*_*target*_, *X*_*lure*_) *< D*_*B*_(*X*_*target*_, *X*_*lure*_) *< D*_*C*_(*X*_*target*_, *X*_*lure*_). Related to mutual information *I*, the Kullback–Leibler (KL) divergence quantifies the dissimilarity between two probability distributions in bits, and information capacity denotes the maximum information a representation can reliably transmit given its signal variance and noise. All three measure, in the currency of bits, how much structure one signal carries about another. The same quantities reappear as the “rate” term in rate–distortion objectives which limits the information a compressed representation may retain about its input.

Compressing redundant information is distinct from how the Marr-Albus expansion model achieves dissimilarity through sparse coding. We can demonstrate this by using *β*-variational autoencoders (VAEs) to compress each stimulus individually *(53, 63, 64)*. In this framework, the KL divergence quantifies the information rate retained by a latent distribution *q*_*ϕ*_(*z*|*x*) relative to the prior *p*(*z*), the reconstruction error quantifies the distortion, and the multiplier *β* controls the degree of lossy compression governed by the rate-distortion function. A trade-off between compression and sparsity emerges in the *β*-VAE’s representations of the task stimuli (**Figure 1F**). This trade-off is also found in biologically-realistic neural network simulations of contrastive Hebbian learning dynamics, which perform dimensionality reduction by converging to the dominant principal components of their inputs (**Figure 1G**) *(71, 83, 84)*. Inputs to this model were synthetic stimuli that we generated to directly control the level of input redundancy and dimensionality as combinations of a predetermined set of latent dimensions. Across both approaches, an increase in sparsity hinders compression, indicative of distinct processes.

The compression and expansion models offer complementary pathways to implement pattern separation by making input representations more dissimilar (**Figure 2A**). In theory, there are countless ways to discard information while increasing the dissimilarity between inputs, including through random signal dropout. Compression constrains this problem to a regime of principled solutions that safely discard redundant information while preserving sufficient fidelity to recognize the original input. For example, targeting the most redundant information (the largest principal components) can enhance memory discrimination by reducing redundancy between similar inputs and isolating their unique, mismatching information. In contrast, discarding information that explains the least shared variance (the smallest principal components) can increase similarity between inputs, consistent with pattern completion (**Figure 2A**).

**Figure 2.**
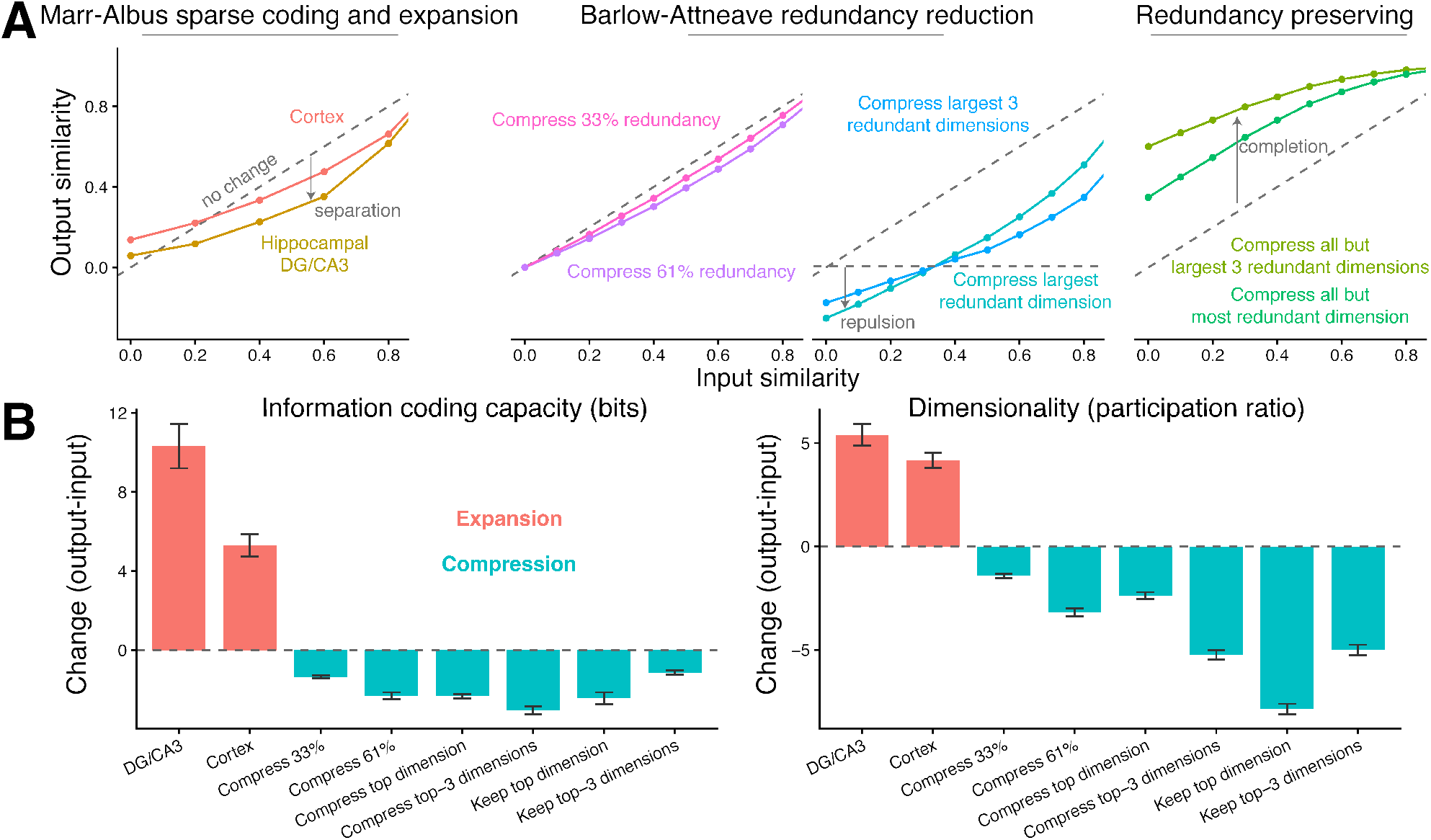
Discriminating between compression and expansion models. Input stimuli are abstracted one-hot representations of categorically structured stimuli, following *(76–78)*. **(A)** *Left*: The canonical pattern separation function, in which similar inputs are transformed into more dissimilar neural representations, replicated based on Norman & O’Reilly (2003) *(13)* in a biologically realistic neural network model of cortico-hippocampal function *(67)*. The hippocampus pattern separates via sparse coding and expanding connectivity to project the input onto separate output neurons according to the established Marr-Albus model. The cortex still pattern separates, albeit more modestly, despite lacking the hippocampus’ sparsity. *Middle*: Barlow-Attneave’s redundancy reduction hypothesis suggests alternative mechanisms for pattern separation based on lossy compression. Some forms of compression can explain repulsion of inputs to be more dissimilar than even unrelated items *(79–81). Right*: Redundancy-preserving compression leads to pattern completion, taking a partial or degraded input and using it to reconstruct a stored memory *(82)*. Different forms of compression can support separation or completion. **(B)** Information theory and dimensionality metrics are diagnostic of either the Marr-Albus sparsity and expansion mechanism or the Barlow-Attneave redundancy reduction mechanism. The Marr–Albus model predicts increased dimensionality and information-coding capacity because its sparse expansion recodes the input onto many more units, distributing the original variance across more dimensions with more uniform importance and amplifying variance relative to a fixed noise floor. The Barlow-Attneave redundancy-reduction hypothesis predicts a more efficient, less redundant code. The code discards the input’s redundant structure across the most frequent, shared units, or the dominant components explaining the most shared variance. We operationalize redundancy reduction as lossy compression, reducing dimensionality and information-coding capacity.

Compression and expansion hypotheses produce discriminable signatures in information theory and dimensionality measures. The compression model predicts information loss and fewer encoded dimensions, whereas the expansion model predicts greater information capacity and more encoded dimensions by distributing information across independent neurons through sparse projections (**Figure 2B**). Together, these models make opposing predictions about how information is allocated and represented in neural codes.

### Modeling image compression from neural network representations

Can the lossiness of compressing inputs explain memory discrimination performance? We calculate the lossiness from the perceptual and semantic feature representations of the stimuli **(Figure 3A)**. Of particular interest are the target-lure pairs of stimuli evoking (1) a single-exposure memory of a target image during the study phase and (2) the subsequent exposure to a similar lure image during the test phase. Analysis-by-synthesis theory posits that perception involves inferring a compact set of latent causes that can reconstruct the input *(85–87)*. Consistent with this view, neural networks represent stimuli as points in an internal space spanned by latent variables that underlie experience *(88)*. Such inference parallels memory reconstruction, where the learned latent structure enables efficient recall. The neural network separately encodes the target and lure as two points, each of which can be decoded back into visual representations.

**Figure 3.**
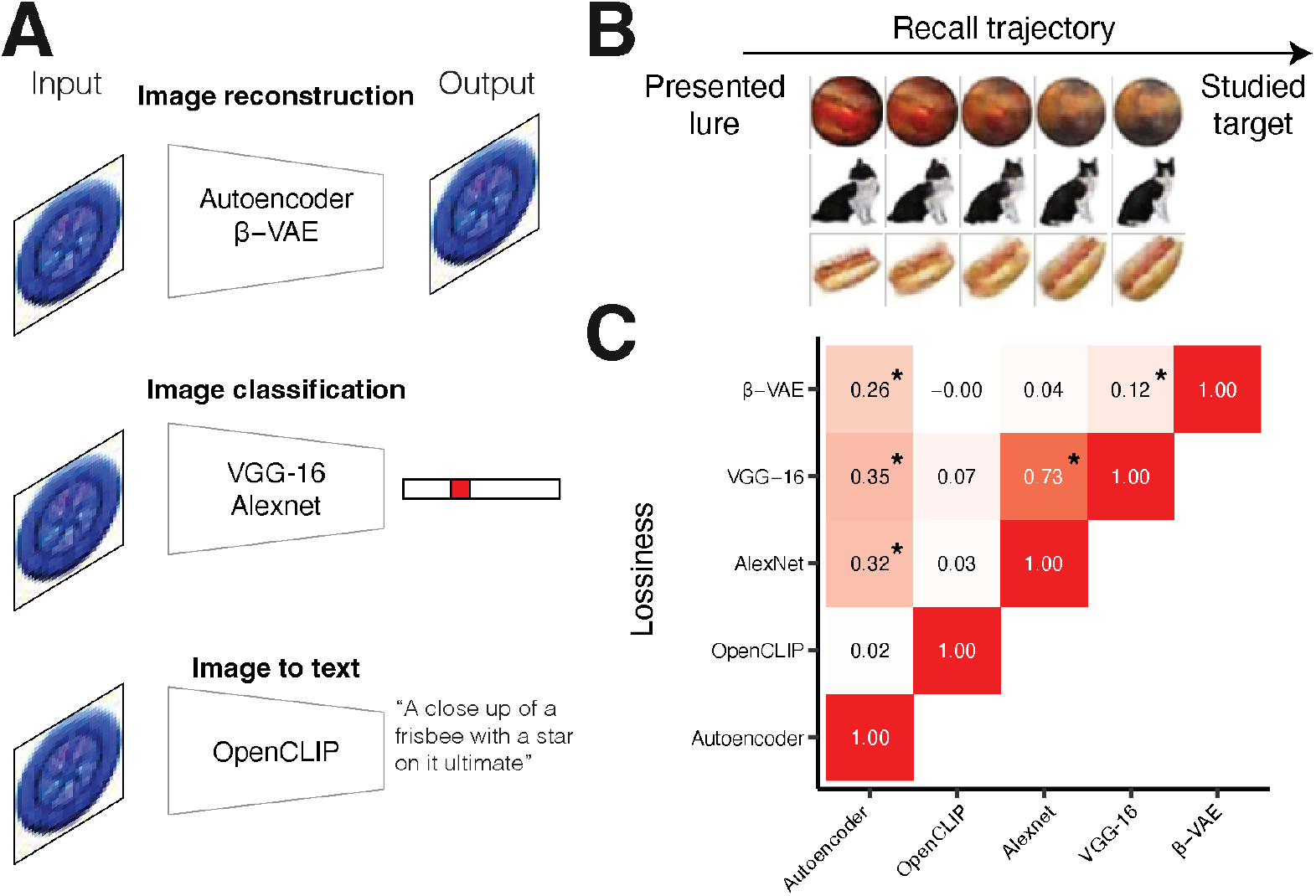
Modeling image compression from neural network representations. **(A)** Neural networks labeled by their respective tasks. Each network was applied to all images *X* to obtain the “sensory” or “semantic” feature vectors *z* used for each task. **(B)** Given the presented lure *z*_*lure*_, the neural network retrieves the studied target *z*_*target*_ by reconstructing a trajectory of representations increasingly similar to the target. To generate the trajectory, a parameter *λ* controls the weighted average of the target and lure representations: *z*(*λ*) = (1 − *λ*)*z*_target_ + *λz*_lure_, where *λ* ∈ [0, 1]. We measure the lossiness of compressing the *z*_*lure*_ to the *z*_*target*_ by applying an information theoretic algorithm to simulated confusion matrices based on the cosine distance between each *z* along a trajectory from *z*_*lure*_ to *z*_*target*_. **(C)** The correlation table shows similarities and differences between the lossiness across models.

Upon viewing the lure as input, we simulate memory retrieval of the target point as the (re)construction of that past experience starting from the lure point. Memory construction occurs via using the neural networks as generative networks *(64, 88)*, which amounts to traversing the space between the lure point and target point by generating representations that had not been experienced using the generative network’s latent variables. This retrieval trajectory generates points that become closer to the target and further from the lure. Decoding the points back to images allows for visual inspection of information content and distortion (**Figure 3B**).

We calculate pairwise compression of redundant information between the target and lure from their retrieval trajectory. Using prior information theoretic methods on the generalization and discriminability of perceptual stimuli *(43)*, we infer the lossiness values which address the question of what lossy encoding would most efficiently account for the observed overlap (i.e., cosine similarity) of the pairs of representations in the network’s retrieval trajectory. It operationalizes the lossiness of the encoding as a cost for encoding one point as another, inferred by searching for the costs under which the observed pattern of overlap would be optimal for a compressor with limited bits of information according to information theory. Low fitted costs are inferred for pairs of points along the trajectory that generative model treats as more similar and redundant.

Although information can be discarded strategically across target-lure pairs, such strategic compression is not possible during memory encoding, when the lure has not yet been encountered. Compressing a single stimulus (i.e., an individual target) may improve the efficiency of its memory representation and store information that explains variation among the stimuli experienced thus far, supporting later discrimination.

However, because encoding cannot anticipate which information will be unique or redundant relative to a future lure, discarding information that later proves distinctive could increase discrimination errors. To consider these scenarios, we investigate how *β*-VAEs compress individual images **(Figure 4A)**. To compare with pairwise compression, we average the lossiness across target-lure pairs. Lossiness values are similar for similar neural networks (e.g. AlexNet and VGG-16, Spearman’s *ρ* = 0.73, *p <* 0.001) and uncorrelated between these models and generative models such as the OpenCLIP image-to-text transformer model (*ρ <* 0.07, *p >* 0.09), suggesting good coverage of convergent and divergent representations of differing sensory and semantic features **(Figure 3C)**.

**Figure 4.**
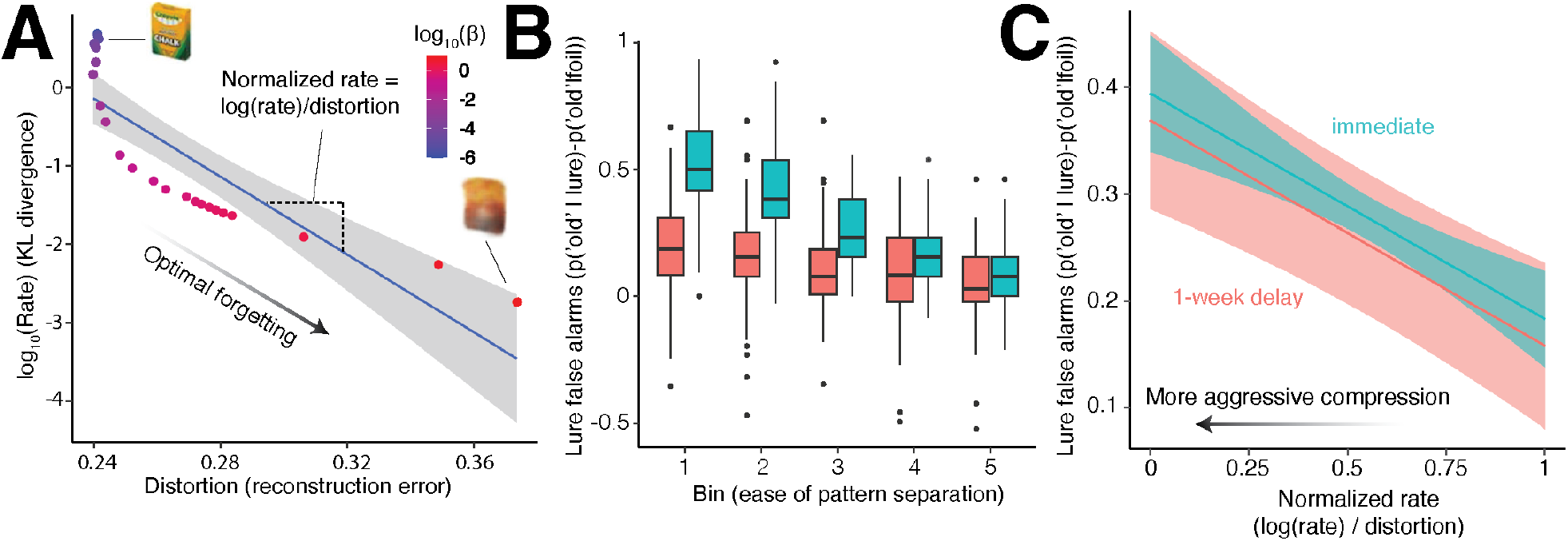
Gist-like memories due to aggressive compression of individual stimuli explains increased false alarms. **(A)** Each individual image undergoes lossy compression in a *β*-VAE, which optimizes the trade-off between information rate and reconstruction error at different values of *β*. Information compression may support memory retrieval and discrimination, but overly aggressive compression can produce gist-like representations that increase false memories (e.g., “a new yellow and green box of crayon chalk in white” versus “a used, crumpled yellow and red bag”). An “optimal forgetting” function *(63)* is quantified for each individual stimulus by the slope of the image’s rate-distortion function, which we term the normalized rate: the amount of information discarded per unit of distortion. Steeper slopes indicate more aggressive compression, whereas shallower slopes indicate more conservative compression. **(B)** Participants’ memory discrimination degrades after 1 week to floor performance. **(C)** Consistent with our hypothesis, participants who viewed images with a lower normalized rate tended to have more lure false alarms in both the immediate and delayed tests. More aggressive compression discards more information for compact gist-like memory representations at the cost of a greater risk of false memory.

### Aggressive compression of individual stimuli creates gist-like memories related to increased false alarms

We first test whether the compressibility of individual stimuli predicts errors in memory discrimination. Errors are quantified by the lure false alarm rate, reflecting confusion of a lure with a studied target. Increasing lossiness corresponds to different levels of information discarded along rate-distortion curves unique to each image, which were estimated using *β*-VAEs. We quantify this difference using the normalized rate per image, the slope of the rate-distortion function, defined as the amount of information discarded per unit of distortion **(Figure 4A)**. More negative, steeper rate-distortion slopes indicate individual images are compressed at a greater cost to fidelity, whereas more positive, shallower slopes indicate more compression is possible without as much detriment to fidelity. We assessed whether the compressibility of individual images predicts memory discrimination errors, as well as when the test phase was delayed by 1 week to introduce greater memory interference and forgetting (**Figure 4B**). Consistent with the compression hypothesis, participants exhibited more lure false alarms for images with lower normalized rates in both immediate and delayed sessions (*b* = −0.21, *p <* 0.001), indicating that aggressive compression can produce efficient but false gist-like memories **(Figure 4)C**. Aggressive compression may therefore generate false memories by limiting the information available to detect mismatches between the studied target and lure.

### Lossy compression of target-lure pairs relates to easier and better memory discrimination

Next, we test whether compression of target-lure pairs supports memory discrimination in the behavioral data. All stimuli are organized into discretized bins of similarity which indicate how easy or difficult they were to discriminate based on prior behavioral data *(69)*. Consistent with the compression hypotheses, we find that target and lure images that are easier to discriminate tend to be those where compressing targets into lures has greater lossiness (all p-values Bonferroni corrected; **Figure 5A**). This effect replicates across two image sets and all perceptual (0.25 *<* Spearman’s *ρ <* 0.40, *p <* 0.001) and semantic models (0.09 *< ρ <* 0.19, *p <* 0.04) that compress targets into lures, but not in *β*-VAE models trained to reconstruct individual items (*ρ <* 0.07, *p* = 1). For pairwise models, lossiness is defined for each target-lure pair, whereas for *β*-VAEs, pairwise lossiness is estimated by averaging the lossiness of the target and lure, each estimated independently to capture information discarded during its individual encoding events. Participants tended to have better memory discrimination of more lossily compressible stimuli (linear regression 0.02 *< β <* 0.38, Bonferroni-corrected *p <* 0.006, **Figure 5B**). This pattern suggests that compressing information within individual stimuli contributes less to memory discrimination, primarily limiting target fidelity, than compressing information that is redundant between targets and lures.

**Figure 5.**
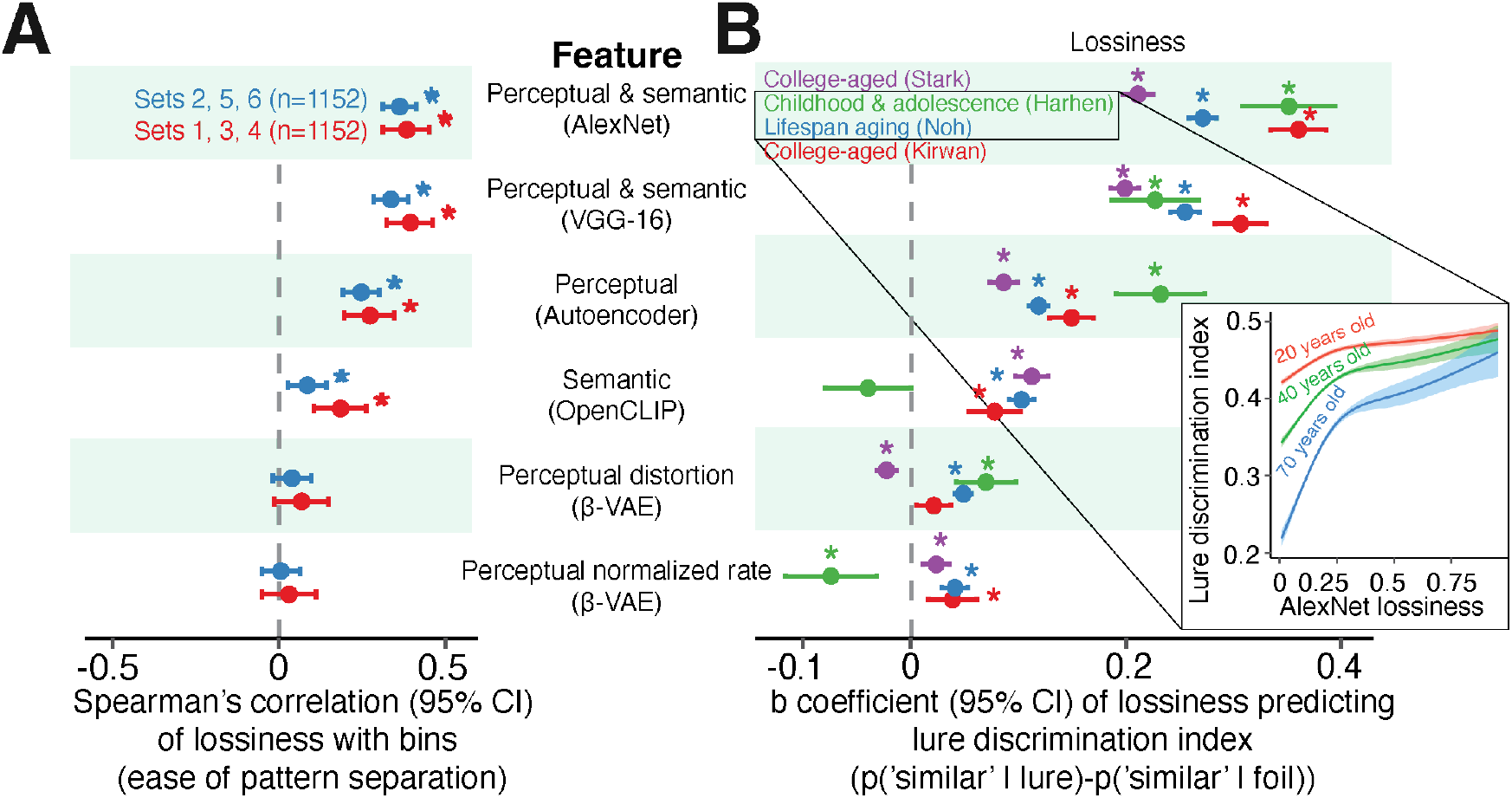
Lossiness of perceptual and semantic compression predicts memory discrimination ease and performance. **(A)** Lossiness is first computed on a stimulus-specific basis then computed on the participant-specific set of stimuli observed as lures or targets. Consistent with our hypothesis, greater lossiness is correlated with easier memory discrimination across two separate image sets. This effect was strongest for perceptual and sensory features. We did not observe an effect of lossiness using the *β*-VAE model. **(B)** Consistent with our hypotheses, greater lossiness is correlated with better memory discrimination across datasets. The effect was greatest for perceptual and semantic features. The effect was less consistent for only semantic features and for the *β*-VAE model. Inset: nonparametric generalized additive modeling was used to flexibly model how lossiness and memory discrimination interact with age. Individuals better pattern separate images with greater lossiness across the lifespan. The steeper logarithmic form for older adults (*F* = 732.1, *p <* 0.001, lossiness-by-age interactions *p <* 0.001) suggests that lossiness may especially help reduce performance gaps between older compared to younger adults.

Next, we assess if age moderates the relationship between lossiness and lure discrimination performance. To this end, we used generalized additive models with penalized splines, a method which allows for statistically rigorous modeling of linear and nonlinear effects while minimizing over-fitting *(89)*. Lure discrimination performance by older adults was more strongly related to the lossiness of compression than younger participants (*F* = 732.1, *p <* 0.001, lossiness-by-age interactions *p <* 0.001; **Figure 5B inset**), consistent with the notion that older adults can use semantic memory to compensate for degradations in episodic memory as their semantic knowledge increases *(90, 91)*. A small amount of lossiness can contribute significantly to improvement, but its benefits exhibit diminishing returns. In support of the compression hypothesis, lossiness predicts memory discrimination ease and performance. Perceptual and semantic features were most related to the ease and performance of memory discrimination.

### Lossily compressed high-level perceptual and semantic representations associated with easier and better memory dicrimination

We next investigated which kinds of image features were most related to memory discrimination ease and performance, comparing low-level features about spatial detail to high-level features about semantic abstraction. In the behavioral datasets, more compression of all perceptual features related to easier memory discrimination (0.26 *< ρ <* 0.40, *p <* 0.001). Higher-order semantic features (representations in deeper layers) had greater effects on the ease of memory discrimination than lower-level features about spatial details (representations in shallower layers; *r* = 0.97, *p <* 0.001; **Figure 6A**). We find a similar relationship when analyzing memory discrimination (0.08 *< β <* 0.83, *p <* 0.001 **Figure 6B**). Participants tended to have better performance when either the deepest or most shallow features were more lossily compressed.

**Figure 6.**
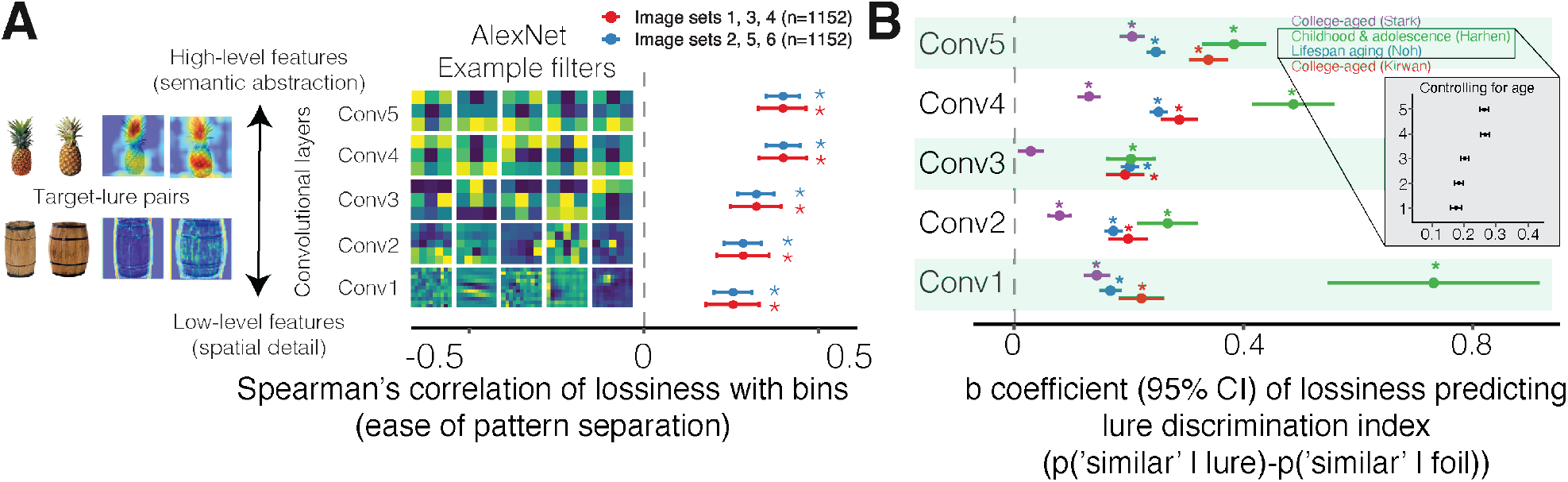
Lossiness of high-level semantic abstractions are more important than low-level spatial details for memory discrimination ease and performance. **(A)** Left inset: AlexNet contains a hierarchy of convolutional layers that process increasingly abstract image features, from low-level perceptual features such as edges, shapes, and textures (fine-grained filters), to higher-level perceptual features that capture conceptual and semantic information for object categorization, and semantic features that represent verbal descriptions of images (coarse-grained filters). Grad-CAM (Gradient-weighted Class Activation Mapping) is an explainable AI technique that highlights the image regions containing information most important for object recognition, allowing us to visually inspect which information is captured by high- and low-level filters *(68)*. Right: Compressions of targets into lures features that incur greater lossiness tended to be better pattern separated. The lossiness of higher-level features characterizing semantic abstraction have a stronger effect on the ease of memory discrimination than low-level features characterizing spatial detail. **(B)** Individual differences in memory discrimination is most strongly predicted by the lowest-level layers for children and adolescents and highest-level layers of processing for older participants. Layers are visualized in descending order by average effect size. Inset: controlling for age results in the higher-level layers being more predictive of performance than lower-level layers.

Whether the deepest or most shallow features are more important may be driven by age. In the dataset of our youngest participants including children and adolescents, performance was more strongly related to low-level sensory details than higher-level features, as they may have not yet formed the richer semantic memories of older adults. To further test whether the order of importance of feature layers is explained by age, we conducted an analysis controlling for age as a covariate. After adjusting for age, the predictive strength of high-level features again surpassed that of low-level features **(6B inset)**, indicating that age may moderate the observed shift from reliance on perceptual to semantic processing (0.21 *< β <* 0.32, *p <* 0.001). This pattern of results suggests that lossily compressing high-level semantic features supports memory discrimination, perhaps characterizing how the ventral visual stream discards inessential information to detect objects *(92–96)*.

### Neural signatures of lossy compression in higher-order visual stream and hippocampus for correct lure discrimination

In light of the computational and behavioral evidence for the compression hypothesis, we next test for compressed neural representations during memory discrimination. We use neural networks to obtain the hypothesized stimulus encodings at different levels of compression and perform representational similarity analysis (RSA). As we had particular interest in the transition from the higher-order end of the visual stream (IT) and the hippocampal region classically implicated in pattern separation (DG/CA3), we use the representations generated in the last convolutional layer of AlexNet, and assess its representational similarity in IT cortex and hippocampal DG/CA3. We hypothesized representational similarity at intermediate compression but not with no compression or overly aggressive compression.

We train a neural network that combines two compression terms in an objective function: a redundancy reduction and an information bottleneck term (**Figure 7A**; see **Materials and Methods**) *(42, 65, 66)*. The network takes as input the AlexNet features of each lure pair and applies a method called Gradient-weighted Class Activation Mapping *(68)* to quantify how important stimulus features are to classify a stimulus’ object category. The difference between the feature importance for lures and pairmates measures the most discriminative features for each pair (a mismatch signal). An encoder is trained to map these discriminative features onto a latent code as a multivariate probabilistic distribution, while a decoder is trained to reconstruct the discriminative features.

**Figure 7.**
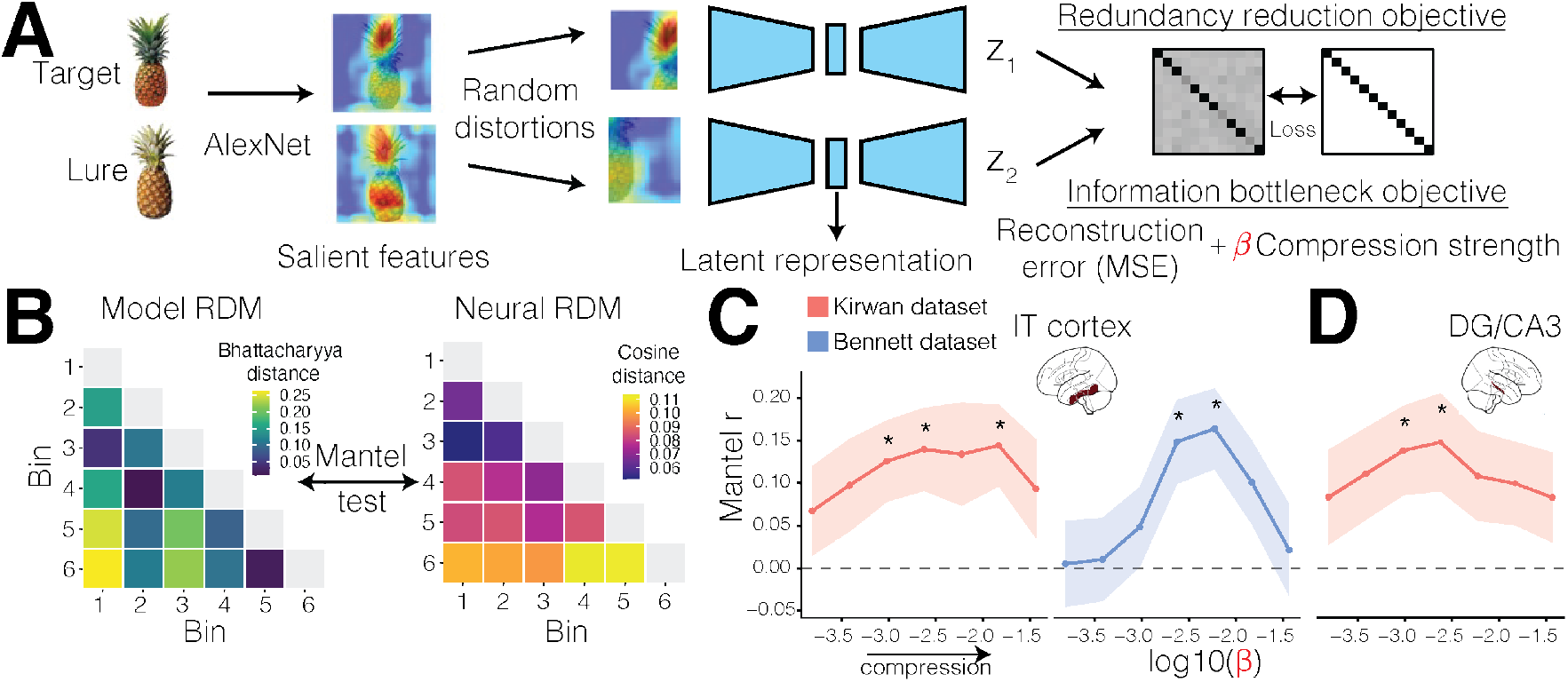
Representational similarity analysis of information bottlenecks in the visual stream and hippocampus. **(A)** Compressed encodings for each lure pair obtained from a neural network model architecture combining redundancy reduction and information bottlneck objectives *(65, 66)*. The parameter *β* controls the strength of compression. The output is the probabilistic discriminability (Bhattacharyya distance) of each lure pair encoding. **(B)** The model-derived distances and neural activity elicited for each lure pair are averaged into the lure pair’s similarity bin, producing representational dissimilarity matrices (RDMs). The representational similarity of model and neural RDMs are compared with a Mantel test. **(C)** In both task fMRI datasets *(37, 38)* and in only lure correct rejection trials, representational similarity of most discriminable object features in IT cortex peaks at intermediate compression. **(D)** In only one dataset and in only lure correct rejection trials, representational similarity of most discriminable object features in DG/CA3 peaks at intermediate compression. *: *p <* 0.05 following FDR-correction for regions and *β*.

The two terms in the objective function govern training. First, the redundancy reduction term in the objective function enforces an orthogonalized code of stimuli. Two distorted views of the same stimulus (random crop, color jitter, flip, and blur) are encoded. The cross-correlation matrix of the two encodings is penalized toward the identity matrix, enforcing invariance of each latent dimension to the distortions and reducing redundancy across dimensions. Second, we evaluate the additional contribution of compression in an information bottleneck term. This term enforces the latent code to have lower reconstruction error and use fewer bits of information used beyond a Gaussian prior. The strength of compression is determined by a multiplier *β* (similar to the *β*-VAE) that amplifies or dampens the penalty on the bits of information used. When *β* approaches 0, the decorrelated representation is effectively uncompressed. Because the compressed codes are probabilistic, we can determine their discriminability as the separation of the multivariate probability distributions measured by the Bhattacharyya distance (a generalized d’ in signal detection theory).

To apply RSA, we average stimulus values within the target-lure similarity bins *(37, 38)*. Model dissimilarity matrices contain the difference in Bhattacharyya distances between compressed codes, whereas neural dissimilarity matrices contain the cosine distance between target and lure activity patterns (**Figure 7**B). Model and neural matrices are compared with a Mantel test at each level of compression strength, with p-values adjusted by FDR-correction across regions and *β*.

Consistent with our hypothesis, representational similarity occurs specifically at intermediate compression. In both the Kirwan and Bennett datasets *(37, 38)*, IT cortex was associated at intermediate compression strengths (**Figure 7C**; Kirwan dataset: Mantel *r* = 0.13-0.14, *p*_FDR_ = .04; Bennett dataset: *r* = 0.15 − 0.16, *p*_FDR_ = .02), with no significant correspondence at the weakest or strongest compression levels tested. DG/CA3 showed the same intermediate-compression profile only in the Kirwan dataset and not the Bennett dataset (**Figure 7D**; *r* = 0.14–0.15, *p*_FDR_ = .04). Experimental control regions M1 and S1 did not reach significance in either dataset nor did they isolate effects to intermediate levels of compression, suggesting these results are not driven by a global or a task-general signal (see **Figure S1**; trending in one dataset with *p* = .06 − .07 at the highest level of compression, but not the other *p* = .81). DG/CA3 correspondence in the Bennett dataset was numerically strongest at the highest compression tested but did not exceed the critical threshold after correction (peak *r* = 0.14, *p*_FDR_ = .09). All correspondences were only observed when analyses were restricted to correct lure rejection trials. When considering all trials, the strongest hypothesis-consistent correspondence was not significant (*p*_*F DR*_ *>* 0.36; control regions *p*_*F DR*_ *>* 0.78). These results are consistent with a compressed code in the higher-order visual stream and mixed evidence for a compressed code in the DG/CA3.

### Theoretical links between retrieval-related states and lossy compression

Different retrieval states during memory discrimination could offer complementary explanations for the putative signatures of compression in the hippocampus. To explore a possible link between retrieval and compression, we investigate memory discrimination in the C-HORSE (Complementary Hippocampal Operations for Representing Statistics and Episodes) model *(13, 67, 77)*. A link between recall and dimensionality reduction could follow from the geometry of Hebbian attractor dynamics *(82, 97)*. In this framework, an *attractor* is a stable pattern of neural activity corresponding to stored memories (e.g., of the studied target stimuli). *Attractor capture* indicates that activity has converged to the attractor point, implementing the retrieval of a stored pattern given a partial or noisy input (e.g., the lure stimuli).

In a recall-to-reject account, exposure to a lure triggers recall of features from its studied target, which are then compared with the lure, and a substantial mismatch supports correct rejection *(13)*. Compression offers one way to amplify this mismatch, by discarding redundant information from the retrieved target. Alternatively, lure discrimination can result from retrieval failure. Pattern separation transforms sufficiently dissimilar lures into representations even more dissimilar from their targets’, so recall is never triggered. Because recall is a high-threshold process that unstudied items rarely elicit, the absence of recall is itself a signal that the lure is distinct *(13, 47)*.

We consider how these retrieval states could be linked to the compression of network activity in two ways. Under an attractor-capture hypothesis, capture concentrates activity: false alarms and recall-to-reject both reflect retrieval-related capture by the target’s attractor and produce stable, low-dimensional, and compressed states. In contrast, retrieval failures (partial completions between attractors, or capture by a wrong neighboring attractor) produce varied, higher-dimensional, uncompressed states. Under a statistical-learning hypothesis, contrastive Hebbian learning converges on the few dominant dimensions of information shared across the studied targets *(71, 83, 84)*. Because stored attractors are pattern-separated, lures captured by distinct target attractors (false alarms) are dispersed across them and are higher-dimensional and uncompressed. In contrast, uncaptured lures (retrieval failures) default to the statistically-learned dynamics of shared target information and settle into mutually similar, lower-dimensional, and compressed states.

C-HORSE networks contain neuronal layers corresponding to entorhinal input, dentate gyrus, CA3, proximal and distal CA1, and neocortex, connected by projections governed by contrastive Hebbian learning (**Figure 8**). We trained 25 networks, each tasked with learning 45 target stimuli (10 feature slots taking one of three values and a localist identity code) to an 80% accuracy criterion. The trained network was then tested on lures of graded similarity to the targets. Each studied target has nine lures of differing similarity (1-9 features altered) and one unrelated foil. Items shared a mean feature overlap of 0.40 with one another. Given the lure’s features, the network was tasked with pattern completing the corresponding studied item’s identity code from the lure’s features. We used a completion threshold of 0.5, such that recall was counted only if at least half of the recalled identity’s code units were active above 0.5 (see **Materials and Methods**). Completions failing this criterion were scored as retrieval failures (recall score of 0) and thereby fell below the decision criterion. The reject decision also occurs if the recall-to-reject score, (match − mismatch)*/*10 features ∈ [−1, 1], fell below the network’s decision criterion, set as the midpoint between the network’s mean recall scores. Discrimination was quantified as *d*^*′*^ at each similarity level and averaged across networks (the z-transformed hit rate over studied targets minus the false-alarm rate over lures). Using these parameters, we first replicate lure discrimination performance and foil correct rejections from prior work (**Figure 8B**), showing that discrimination decreases with greater target–lure similarity *(13)*. We also replicate the pattern-separation function across stimuli and networks (**Figure 8C**).

**Figure 8.**
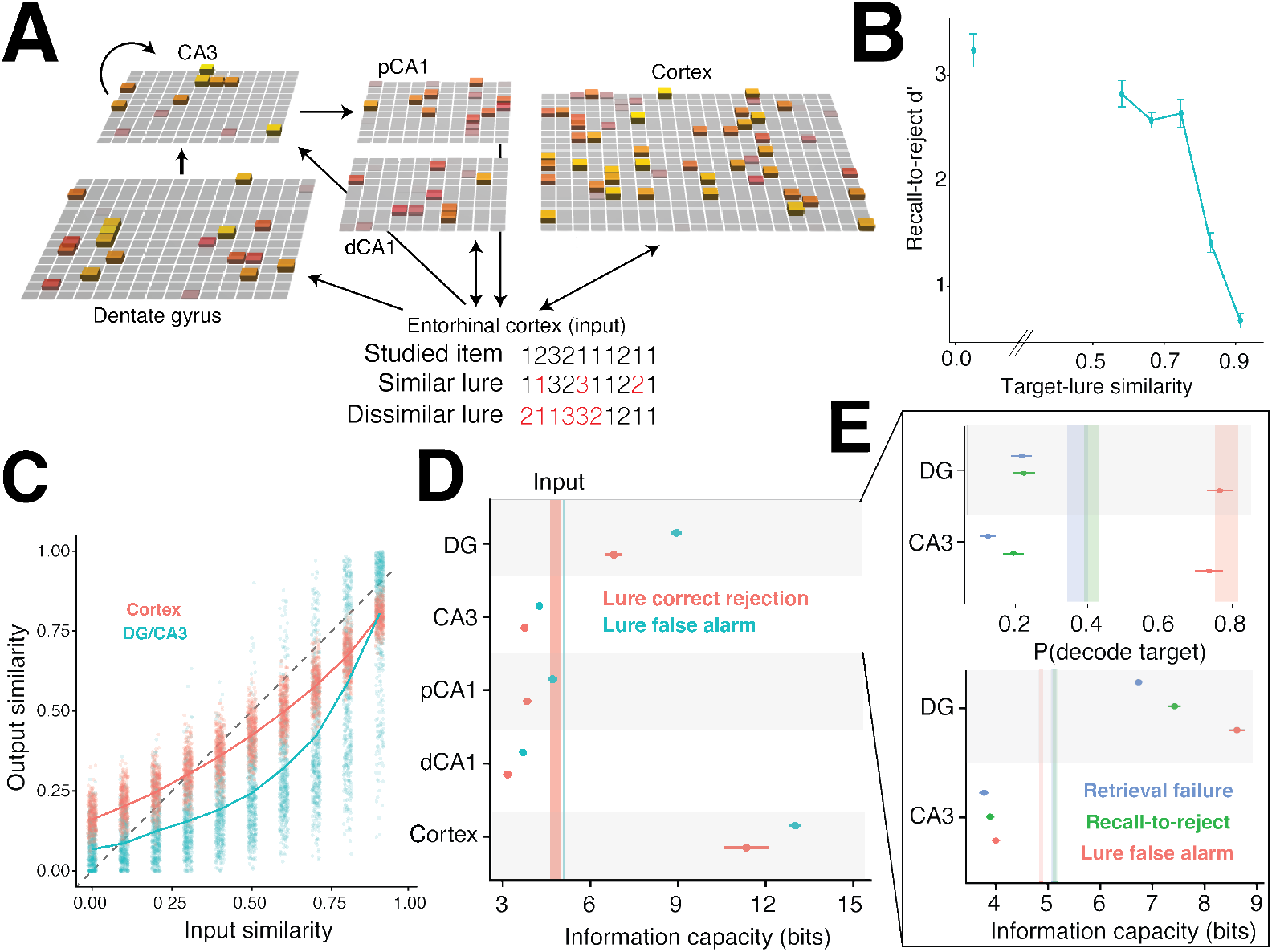
Retrieval-related compression and expansion in a cortico-hippocampal neural network. **(A)** A biologically realistic cortico-hippocampal model with sparse neural activity (column height depicts magnitude) *(67)*. Synthetic target and lure stimuli were generated with differing levels of similarity. **(B)** Memory discrimination performance based on hippocampal retrieval, replicating model behavior in Norman & O’Reilly (2003) *(13)*. Performance worsens with greater similarity. **(C)** Pattern separation function across all networks and stimuli, with the hippocampus implementing Marr-Albus expansion and sparse coding. **(D)** Information capacity exhibits the same pattern across all regions, where lure correct rejections (red) are reduced compared to lure false alarms (blue). Dimensionality reduction follows same pattern as information capacity loss (not shown in figure due to redundancy). Dentate gyrus (DG) and cortex expand relative to the input baseline (shaded vertical bars whose widths indicate 95% CIs), whereas auto-associative CA3 and hetero-associative CA1 regions compress. **(E)** For DG and CA3, we disambiguate the correct rejection trials by whether they arose from recall-to-reject and retrieval failure pathways. Consistent with its qualitative mechanism, retrieval failure reflects the most information loss as a corresponding target is unable to be identified. Lure false alarms exhibit the most information retained since there is a high probability of mistaking the lure by decoding it as the target. Recall-to-reject shows an intermediate level relative to the other pathways. Dimensionality reduction follows the same pattern as information loss (not shown in figure due to redundancy). The input baselines are depicted as shaded vertical bars whose widths indicate 95% CIs.

To investigate compression we measured each region’s dimensionality and information capacity using the activation covariance across lure trials (the metrics that we showed discriminate between compression and expansion in **Figure 2**). For each lure similarity, we drew a correct-rejection sample with the same total sample size as the false-alarm group to control for sample size imbalance. Dimensionality and information capacity was computed on each resample and averaged over 100 resamples. We observed loss of information capacity and dimensionality reduction for correctly rejected lures compared to falsely recognized lures, comparing equal numbers of trials at the same target-lure similarities (**Figure 8**D). Auto-associative CA3 and hetero-associative CA1 regions compress inputs relative to baseline, consistent with proposed roles for CA3 in compressed attractor-based retrieval and CA1 in detecting mismatches between inputs and DG/CA3 outputs *(97–100)*. Correctly rejected lures loaded more of their variance on the three leading shared components of activity than false alarms in DG (Wilcoxon signed-rank test *p <* 0.001) and CA3 (*p* = 0.001) with no difference in the cortex, consistent with the statistical learning hypothesis.

Disambiguating the correct rejections into recall-to-reject and retrieval failure pathways, we compared regional decodability, dimensionality, and information capacity relative to the input statistics of the EC as a baseline (**Figure 8**E). Target-identifying information was always reduced relative to the input, consistent with the data processing inequality. However, falsely recognized lures preserved it almost fully (Δdecodability = −0.02 in DG, −0.05 in CA3), whereas both rejection routes lost more information (−0.16 to −0.25). Successful completion of the target preserves the input’s identity information, and correct rejection discards much of it. The change in capacity and dimensionality revealed that DG expands while CA3 compresses relative to the input, due to the regions’ inherent circuit properties (i.e., sparsity and size). For memory discrimination, these regions’ activity exhibit a consistent ordering such that false alarms carried the most input-referenced capacity and dimensionality, retrieval failures the least, and recall-to-reject rejections in between. Correct rejection is thus accompanied by graded lossy compression relative to the input, strongest for retrieval failures, independent of whether the region’s overall transform is expansive (DG) or compressive (CA3). These measures are consistent with the statistical learning hypothesis: false alarms decode to distinct stored targets and are dispersed across them (higher dimensionality), whereas correctly rejected states are mutually similar and concentrated on the targets’ shared information (lower dimensionality). Taken together, retrieval-related states could explain observations of low-dimensional, compressed activity in the hippocampus.

## Discussion

In summary, we found behavioral, computational, and neural evidence supporting a lossy compression account of memory discrimination. To support memory discrimination, the cortex may perform efficient compression, whereas the hippocampus may preserve information through sparse expansion and compress it during retrieval.

### Novel theoretical contributions

In contrast to Marr-Albus sparse expansion, the Barlow-Attneave compression hypothesis has the potential to explain the role brain regions that have been implicated in pattern separation, yet lack the hippocampus’ distinctive expansion or sparsity mechanisms *(13, 15, 23, 35–41)*. The compression hypothesis extends the efficient coding principle, where the brain maximizes the amount of information transmitted in an economical form by reducing redundancy *(42, 101)*. This coding strategy allocates limited resources to where they are most relevant for the task at hand *(42)*. In memory discrimination tasks, the most relevant information to encode is the distinctive, mismatching information between target-lure pairs.

The idea that redundant information should be partitioned from unique information is not new. Contrast theory’s distinction between common versus distinctive feature sets *(102)*, recall-to-reject’s split between prototype and item-specific feature slots *(13)*, perceptual-mnemonic accounts of feature conjunctions *(103–105)*, predictive processing accounts of expected and unexpected information *(10, 106)*, and selective attention to shared versus unique features *(107)* all share a common principle: discard redundant features to minimize interference between inputs. Our novel contribution is to unify these ideas under the computational framework of compression, a process that optimizes for redundancy reduction while retaining enough fidelity to support other tasks such as object recognition.

Several theories propose that the hippocampus compresses information through mechanisms analogous to autoencoders, which perform nonlinear dimensionality reduction, or VAEs, which formalize compression through rate-distortion theory *(81, 108–111)*. Autoassociative CA3 may act as a compressive, attractor-based retriever, while CA1 may support the selective compression of expected information through mismatch or prediction-error detection. *(98–100, 106)*. The present work highlights, however, that hippocampal compression is fundamentally constrained by the sparsity mechanisms already known to support hippocampal pattern separation, yet may nevertheless arise during retrieval.

### Computational advantages and disadvantages of compression and expansion

Both compression and expansion can be computationally beneficial for cortico-hippocampal circuits, where the cortex performs efficient compression while the hippocampus implements information-preserving sparse expansion and compressed retrieval *(5, 34, 45, 62, 112–122)*. Switching between differing levels of expansion and reduction may depend on dynamically altering inhibition-induced sparsity *(123, 124)*.

Marr-Albus sparse expansion is energetically efficient *(13, 125, 126)*, can improve the storage capacity of associative memories (though see *(127)*), and can reveal the dimensional structure within information in a form that facilitates downstream processing *(128)*. However, sparsity can also amplify noise, make representations oversensitive to inconsequential input differences, and make it harder for downstream regions to detect the few active units among many.

In parallel, compression via Barlow-Attneave redundancy reduction may support energetic efficiency *(42, 45, 101, 129)* and be most beneficial when the input is high-dimensional and contains redundant statistical regularities. However, it is unclear how much redundancy is truly compressible *(130)*. This remains challenging not only due to differing degrees of redundancy but because seemingly irrelevant redundancy may be retained deliberately for stability and error-correction *(45, 119, 131, 132)*. In contrast to the well-established account of sparse expansion in Marr–Albus theory, how the brain implements information compression remains unclear. Finding low-dimensional structure in neural recordings shows that activity is highly compressible, but more work is needed to demonstrate if the brain is actively compressing information *(45, 118, 133, 134)*.

### Brain-wide pathways to compression

In the present work, we specifically aimed to pinpoint compression and expansion mechanisms in the IT cortex of the higher-order visual stream and in the DG/CA3 of the hippocampus, regions previously implicated in object recognition and pattern separation *(14, 37, 38, 69, 93)*. Other cortical regions are also implicated in compression, such as prefrontal regions associated with selective attention *(44)*. Selective attention may support perceptual and memory discrimination by isolating diagnostic information *(44, 135–139)*. Prior work suggests that even earlier visual regions may perform simlar functions. Serre et al. (2007a, 2007b) *(112, 113)* developed a brain-inspired model of object recognition that alternates between compression and expansion. Early visual cortex V1 has a degree of sparsity which supports known properties of simple cells’ receptive fields, and this sparsity can already begin to expand and decorrelate perceptual inputs *(125, 130, 140)*. Moving to yet earlier stages of visual processing, the lateral geniculate nucleus of the thalamus may also have functional relevance to compression and efficient coding *(141)*. Beyond isolated regions or circuits, compression can be mathematically modeled as a function of network communication across the whole brain *(45, 118)* or as the consequence of Hebbian learning dynamics that converge to representations of the largest dimensions of activity *(22, 71, 83, 84)*.

### Compressible information is task-dependent and changes with age

Different types of information can be deemed compressible or forgettable according to what information is task-relevant. In memory discrimination tasks, we found that high-level information (e.g., parts) was more important than low-level information (e.g., edges, orientations, and textures). This relative importance may change with aging *(91)*. We observed that young participants’ performance was uniquely related to the lossiness of the lowest-level features, a relationship that was moderated with age. Memory discrimination in children may be supported by the neurodevelopment of high-resolution memories for perceptual details *(142–144)*, while loss of precision in memory representations is related to episodic memory decline during aging *(145)*. Older adults’ performance was closer to that of younger adults when the stimuli had lossier compression of perceptual and semantic features. These results are consistent with the notion that the development of semantic knowledge may help organize stimuli in memory, providing a scaffold for perceptual memory encoding *(52, 137, 146–151)*.

### Compression signatures may reflect multiple mechanisms

Putative signatures of compression can be explained by operations that do not necessarily require only compression. For example, reduced dimensionality could also arise from retrieval and predictive processing frameworks *(10, 88, 152, 153)*. The C-HORSE model can decompose each lure into the component predicted by the network’s learned weights and the unpredicted component. The unpredicted component grows with greater target-lure dissimilarity, characteristic of a novelty signal that could be used to discriminate lures.

### Strengths and limitations

This work has several strengths and limitations. First, lossiness was inferred from an information theoretic method developed to fit empirical data from perceptual identification experiments *(43)*. Here, we adapted the fitting procedure to the memory discrimination task, identifying a compression scheme sufficient to generate the observed dissimilarity; but it does not establish that compression, rather than the similarity structure itself, is the generative cause. This concern is partly addressed by using alternative VAE methods for estimating lossiness, which estimate compression from the representations directly rather than from their similarity structure. However, when analyzing *β*-VAE representations, one dataset *(50)* had statistically significant correlations in the unhypothesized direction versus the three other datasets in the hypothesized direction (**Figure S3**). This could reflect a known limitation of VAEs in compressing photorealistic images *(63)* or suggest that compressing stimuli individually (versus pairwise) does not strongly support memory discrimination.

Second, our results directly modeled representations of photorealistic, everyday objects. These stimuli are higher-dimensional (e.g., sharpness, position, form, color, size, pattern, state, usability, function, number of items, and composition) *(137)* than previously considered datasets in visual memory studies applying compression *(63, 64, 88)*. Thus, our work is a step towards a more general understanding of memory discrimination and compression in naturalistic settings. This work demonstrates the practical utility of applying computer vision models as tools to obtain reconstructible and interpretable memory representations. However, the manner in which the neural networks arrive at those representations is unlike human vision *(154)*. This motivated our investigation of biologically realistic models of hippocampal and cortical function during memory discrimination *(30, 71)*, but the trade-off is that such models currently abstract away visual content into nominal indexes of a sparse dictionary code which cannot be used to reconstruct the original visual information contents.

Third, our analysis is limited by the spatiotemporal resolution of fMRI data and by single-trial exposure to each stimulus. For example, dentate gyrus and CA3 cannot be reliably separated at this resolution. We addressed the latter limitation by averaging neural responses within stimulus similarity levels in the RSA and by obtaining convergent results from a biologically realistic cortico-hippocampal model *(13, 67, 77)*. Future task designs using fMRI measures of compression could further minimize stimulus-irrelevant activity through repeated exposure or repeating stimuli within categories *(106, 155–158)*.

## 3 Materials and Methods

### Experimental task

#### Mnemonic similarity task

Participants were instructed to judge object images as indoor or outdoor during 128 study trials (**Figure 1B**). This study phase is intended to have participants incidentally encode the stimuli in memory while performing the indoor and outdoor cover task. Next, during a test phase, participants judge stimuli as “new,” “similar,” or “old,” when the presented stimulus was already seen, never seen, or slightly altered (repeat, foil, and lure trials, respectively) over 192 test trials. Several performance metrics can be calculated that index different memory processes. Memory discrimination performance is measured as the lure discrimination index, or the proportion of correct rejections adjusted by a response bias: *p*(“similar”|lure) − *p*(“similar”|foil). The lure discrimination index in an independent sample was used to discretize the “mnemonic dissimilarity” of original and lure images into 5 bins of ease, where 5 is the easiest and most dissimilar and 1 is the hardest and most similar (**Figure 1C**). A recognition score is defined as correct detections corrected by the false alarm rate: *p*(“old”|repeat) − *p*(“old”|foil). We primarily use the recognition score for quality control purposes, as we focus here on pattern separation whereas this score centers pattern completion processes *(159)*. Finally, lure false alarms are defined as as the proportion of trials where lures were mistaken as a repeat corrected for a response bias: *p*(“old”|lure)−*p*(“old”|foil). This task allows us to probe pattern separation ease, performance, and errors. The stimulus images were obtained from Photodisc, Inc. and use of the low-resolution images here was granted without any restrictions.

#### Data and pre-processing

We analyzed four behavioral datasets. All participants gave written informed consent prior to data collection. Two datasets covered a wide range of ages: a sample of participants across the lifespan (*n* =366; 46 ± 19 years old; 218 women, 144 men, 4 other) *(49)* and a childhood and adolescent sample (*n* =92; 16 ± 5 years old; 54 women, 37 men). The lifespan dataset was recruited from Amazon Mechanical Turk. Child and adolescent data was recruited from Hartley Lab Participant database at New York University. Participants were recruited from the Hartley lab database which includes individuals recruited through ads on social media (e.g., Facebook and Instagram), word of mouth, local science fairs, and flyers on New York University’s campus. For participants under 18 years of age, informed consent was first obtained from a parent or legal guardian. For participants younger than 12 years, a parent/guardian then read an age-appropriate assent form aloud to the child, after which the child provided verbal assent to participate. Participants ages 12 to 17 read an age-appropriate consent form and provided their consent before beginning the study. Participants who were 18 years of age or older provided informed consent on their own behalf. Study procedures began only after the appropriate parental permission and participant assent or consent had been obtained. Participants who had not previously completed an in-person study with the lab completed a brief Zoom call with a researcher. During this call, participants (and their parent or guardian, if the potential participant is under 18 years) were required to be on camera and confirm their full name and date of birth. Adult participants and parents of child and adolescent participants were additionally required to show photo identification.

We also assessed two datasets of undergraduates recruited in a university setting: a cross-sectional sample of undergraduate students who participated for course credit (*n* =208) *(50)* and a longitudinal sample (*n* =78) collected immediately and after 1 week *(51)*. The cross-sectional sample was collected at the University of California - Irvine, and the longitudinal sample was collected at Brigham Young University. Participants in the longitudinal sample completed two study and two test phases, counterbalanced across two sessions. Half of the participants completed the test immediately after study, followed by the second study phase in the same session. They then took their second test one week later. The other half of the participants completed only the study in the first session and then completed the test one week after. They then completed the second study and second test during the second session.

We performed several quality control steps to arrive at our final sample. First, we performed trial-level quality control. If a response was faster than 300 milliseconds or slower than 3 seconds, then the trial was excluded. Next, we performed participant-level quality control. If the participant missed more than 20% of trials, the participant was excluded. Participants also were required to meet minimum scores on the lure discrimination index and recognition metrics indicating minimal engagement with the task. A minimum lure discrimination index of 0 was required; this minimum occurs when response bias was equal to correct rejections indicating chance-level performance. A minimum recognition score of 0.5 was required; this calculated as the correct detection minus a response bias, where chance performance is again 0. For the longitudinal dataset, we only applied the performance thresholds to exclude participants based on their immediate test.

#### fMRI mnemonic similarity task

We analyzed two independent fMRI datasets to test the replicability of findings. The Kirwan dataset was collected at Brigham Young University and was included in a prior publication *(37)*. Participants gave written informed consent prior to completing pre-scan questionnaires and scanning. Participants (n=48; 22.9 ± 3.6 years old, 27 female, 21 male) completed a continuous recognition version of the mnemonic discrimination task *(26, 160)*; see previous publications using this dataset for more details *(37)*. A series of images of everyday objects were shown in sequence. The image set differs and predates those from those published in the prior online implementations of the task, which were used in our behavioral data above *(161)*. Participants judged whether each image was new (foil trials), similar (lure trials), or old (repeat trials). The delay between the first and repeated presentation of lure or repeated objects varied with a mean lag of 19 trials. Six blocks of 107 stimuli (total of 642 images), were presented. Each block contained 32 first presentations, 16 targets, 16 similar lures, and 43 unrelated foils. Participants were required to respond within 3.0 s, after which time the stimulus was replaced by a blank screen with a fixation cross for .5 s followed by the next stimulus.

The Bennett dataset was collected at the University of California - Riverside and was included in a prior publication *(38)*. Participants (*n* = 54, mean age 46.6 ± 27 years, 28 female) provided written informed prior to completing data collection. During fMRI scanning, participants completed a study phase consisting of a sequence of 128 target objects and a test phase consisting of 192 probe objects that were either exact repetitions of objects presented in the study phase (64 repeated targets), similar to objects presented in the study phase (64 similar lures), or novel first presentations (64 novel foils). While the Kirwan dataset employed a 3-choice response format (“old,” “similar,” “new”), this task used a 2-choice format (“old”, “new”). All stimuli were presented on a white background for 2.5 s with a 0.5-s inter-stimulus interval.

### Neural network feature representations

We aimed to use neural networks as tools to operationalize the psychological dichotomies of perceptual and semantic memory. For example, deep-layer representations in Alexnet are referred to as more “semantic”-like because AlexNet undergoes supervised training on a broad array of object categories. These features are referred to as semantic in sense that they represent nameable facts and properties about each class, accumulated across a larger set of experiences akin to the accumulation of semantic memory *(58)*. For shallow-layer representations or the hidden layers of the smaller networks (autoencoder and variational autoencoder) produced via unsupervised learning on less data, we refer to these features as more “perceptual”-like because their size and training diet limits their capacity to learn patterns beyond the lower-level visual information of stimuli.

The perceptual and semantic labels can be inconsistent with the neural networks’ capabilities. For example, it is possible in principle for architectures such as the variational autoencoder to learn semantically interpretable latent dimensions underlying stimuli when trained on many examples and a curated dataset with many exemplars to tile the latent space, but we did not find that this was the case for our experimental stimuli when visually inspecting the decoded representations in the latent space.

All models were implemented using PyTorch. Autoencoder models were trained on 2,252 images across 6 image sets used in the mnemonic discrimination task. Images were originally 400×400 (x3 RGB color channels) and were downsampled to rescale according to smaller input dimensions required by the first layer of each neural network and to reduce computational costs of training. Images were normalized by the mean RGB value across all images to stabilize optimization and improves generalization. Image classification and image to text models were pre-trained on large datasets, described below. Given a stimulus from the mnemonic similarity task, the activations of the hidden layer in the autoencoders were used as the learned feature representation per image, while the activation of the convolutional or fully connected layers were used as feature representations or low-level to high-level perceptual and semantic features in the pre-trained models. The dimensionality of the layers are described in more detail below.

#### Image reconstruction

We use a convolutional autoencoder to learn compressed memory representations of color images in an undercomplete latent space. Stimuli were rescaled to 32×32 (x3 RGB color channels) and pixel intensities were normalized to the range [−1, 1]. The encoder progressively reduces the input image to a compact latent representation using five convolutional layers (three of which use stride-2 downsampling, interleaved with two stride-1 layers; 32 base channels, doubling to 64 in deeper layers) with GELU activations. The tensor produced by the last convolutional layer is flattened and passed through a fully connected bottleneck layer projecting it to a 256-dimensional latent space. The decoder first maps the latent vector back to a 4 × 4 feature map via a fully connected layer, then uses transposed convolutions to upsample the latent representation back to the original spatial dimensions, with GELU activations and a final Tanh activation to bound the output to match the input normalization. We used a reconstruction loss defined as the squared error summed over all pixels and channels of each image and averaged across the batch, and trained for 250 epochs with a batch size of 32 using the Adam optimizer with an initial learning rate of 0.001, reduced by a factor of 0.2 (to a minimum of 5 × 10^*−*5^) when validation loss plateaued for 20 epochs. The dataset was randomly partitioned into training (80%) and validation (20%) sets.

We also use a *β*-VAE to learn a compressed latent representation *z* of the input images *X (53, 162)*. In contrast to the autoencoder above, which maps each image to a single latent point, the *β*-VAE encodes images as distributions over latent space and adds an information theoretic penalty that regularizes the latent space toward a Gaussian prior. These neural networks learn a probabilistic generative model showing how the input data *X* depends on unobserved latent variables *z* and approximate the optimal posterior distribution *q*_*ϕ*_(*z*|*x*) over the latent factors given the observations. The objective of a VAE is to minimize the distance (KL-divergence) between the approximate *q*_*ϕ*_(*z*|*x*) and true posterior distribution *p*(*z*|*x*) by maximizing the evidence lower bound, described in the next section. The latent variables *z* can also be used to reconstruct inputs *X* with a measurable error. The KL-divergence and reconstruction error term in combination forms the training objective of a variant of the VAE called the *β*-VAE: minimizing reconstruction error (binary cross-entropy) and minimizing the KL-divergence term to varying degrees according to the scalar parameter *β*. The objective function is:

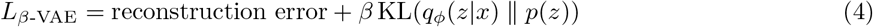

Increasing *β* enforces stronger constraints on the KL-divergence; in practice, this results in greater regularization of the latent space. In sparse *β*-VAEs, we further constrain the loss function to penalize a L1 regularization term on the magnitude of latent representations *z*.

The encoder consisted of four convolutional layers with ReLU activations, progressively reducing spatial dimensions while increasing feature depth. A fully connected bottleneck layer *z* with 512 dimensions parameterized the mean and covariance of the latent distribution, and a latent vector was sampled using the reparameterization trick using a Gaussian distribution parameterized by *ϕ*. The bottleneck layer factorizes the distribution over inputs *X*, forming a probabilistic representation that can disentangle the primary factors of variation across observations. The decoder, composed of transposed convolutional layers with ReLU activations, reconstructed images from the latent space. The final layer of the decoder applied a sigmoid activation to ensure output pixel values remained between 0 and 1. We trained the models over 5 random seeds using a batch size of 128 over 200 epochs with the Adam optimizer with a learning rate of 0.001. Each of the 5 seeds performed a sweep across 22 *β* values ranging from 10^*−*6^ to 10 to estimate the rate-distortion function of each image.

#### Image classification

We used two pre-trained convolutional neural networks called AlexNet and VGG-16 to encode different feature representations of the task stimuli. We refer to these representations as perceptual and semantic features. Stimuli were rescaled to 224×224 (x3 RGB color channels). Both networks were trained on a large number of images across a diverse set of categories in the ImageNet dataset, consisting of 1,281,167 training, 50,000 validation, and 100,000 test images across 1,000 semantic categories. The networks were trained to categorize images into the 1000 semantic categories.

AlexNet consists of nine layers. Briefly, the input layer maps unit activation to colors. Next, the output is processed sequentially by five convolutional layers, containing spatial filters that convolve with the input from the previous layer to produce the 3D tensor representing spatial activations of a particular visual pattern (see **Figure 6A**). Similar to the brain’s ventral visual pathway, the hierarchical structure of convolutional layers are thought to extract low-level spatial details first then more and more abstract higher-level semantic representations in each subsequent layer to support image categorization. Next, there are three fully connected layers which flatten the 3D tensor into a vector.

For each image in the mnemonic similarity task, we obtained six vectors which we refer to as perceptual and semantic feature representations. We obtained a 4096-dimensional vector from the penultimate fully connected layer that the following layer uses for semantic categorization. The penultimate layer is used instead of the last fully connected layer in order to obtain more generalizable, less task-specific, and richer features that are useful for transfer learning, clustering, and similarity tasks. In contrast, the last layer’s features are tailored for categorizing images into 1,000 categories. We also obtained five feature representations with differing dimensionality from the activation of each convolutional layer. The feature representations were 96-dimensional in the first layer, 256-dimensional in the second layer, 384-dimensional in the third layer, 384-dimensional in the fourth layer, and 256-dimensional in the fifth layer.

VGG-16 is another convolutional neural network trained to categorize images on the ImageNet dataset but has a different architecture. The input dimensionality for images is again 224×224×3 and the network is again trained to classify images into 1000 categories. The input image is processed by 13 convolutional layers and 3 fully connected layers. For each image in the mnemonic similarity task, we obtained a 4096-dimensional feature representation from the penultimate fully connected layer.

#### Image to text

To obtain semantic feature representations, we used the contrastive language-image pre-training (CLIP) neural network framework with the ViT-H/14 image embedding variant, a hierarchical vision transformer. This model was pre-trained with the LAION-2B English dataset, a dataset of 2 billion English image-text pairs. Briefly, a vision encoder transforms images into vector representations, using pixel patches. The text encoder processes text descriptions through a transformer architecture, creating word-vector embeddings in the same semantic space as the visual features. The model is trained to optimize a contrastive objective, maximizing similarity between paired image-text embeddings while minimizing similarity with unpaired samples. This objective constrains the vision and text encoders to learn representations in a joint embedding space where semantically similar concepts are more similarly represented regardless of whether their source was an image or text. The resulting image embeddings from this model can be used for zero-shot image classification and other tasks without requiring task-specific fine-tuning. For example, the model performs zero-shot categorization of the Imagenet dataset with a 78.0% accuracy despite not being explicitly trained to do so. For each image stimulus in the mnemonic similarity task, we use this pre-trained model to obtain a text description. This description is tokenized and inputted to the pre-trained text encoder to extract a 768-dimensional word-vector embedding. We refer to the 768-dimensional vectors as semantic features.

### C-HORSE architecture

Simulations used the C-HORSE architecture *(67, 77)*, comprising hippocampal and cortical pathways. The input consists of ten feature layers (one per stimulus slot, each a one-hot code over its three values), a category layer, and a localist identity layer (CodeName; a 6 × 45 grid, one six-unit column per stimulus). These input is considered to serve as the model’s entorhinal cortex interface with the hippocampus. The entorhinal cortex in our analyses denotes the clamped input features, whose activity is the stimulus itself. The hippocampal pathway comprises the dentate gyrus (DG; 15 × 15 units), CA3 (12 × 12), and proximal and distal CA1 (pCA1, dCA1; 10 × 10 each). The cortical layer (CTX) contains 20 × 20 units.

The trisynaptic pathway processes inputs to the DG, CA3, pCA1, and back to the input. These subregions are connected by sparse random perforant-path projections to DG (each DG unit receiving from a random 60% of input units) and to CA3 (10% connectivity), mossy-fiber projections from DG to CA3 (10% connectivity), recurrent CA3 collaterals, and all-to-all (fully connected) CA3→pCA1 and pCA1→input projections. dCA1 is reciprocally connected to the input layers (the monosynaptic pathway), as is CTX (the cortical pathway). All projections have modifiable weights except the mossy fibers, which are fixed at strong values (initial weight 0.9, relative weight scale 10, with the recurrent collaterals at scale 2). They act as a strong, unchanging relay that imposes DG’s sparse, pattern-separated code on CA3. The modifiable projections learn by contrastive Hebbian learning with the pure-Hebbian component disabled (CHL, error-driven). Hippocampal learning rates were fast (0.05–0.1) and cortical learning rates slow (0.0001), so the hippocampus learns the specific studied items rapidly while the cortex accumulates structure slowly. Each of the 25 networks was initialized independently, such that the sparse projections’ connectivity patterns (i.e., which units connect) and all projections’ initial weight values were drawn at random per network.

Within each layer, activity was constrained by k-winners-take-all (kWTA) inhibition: a feedback inhibition level is computed each settling cycle such that only approximately the *k* most strongly driven units remain active, approximating the effect of pooled inhibitory interneurons *(71)*. Inhibition was strongest in DG and CA3 (*G*_*i*_ = 2.8 and 5.0, versus 2.1-2.6 elsewhere), yielding sparse hippocampal codes (∼7% of DG and ∼5% of CA3 units active per pattern, versus ∼8–10% in CA1 and cortex).

#### Stimuli

The training stimulus set contained 45 target stimuli, each defined by 10 feature slots taking one of three values. All targets were variants of a single prototype: mean feature overlap was 0.55 with the prototype and 0.40 between targets. Each target was additionally assigned a unique identity code (one column of the CodeName layer) and a common category unit.

The test stimulus set contained 450 probes. For each target, nine lure probes were constructed with graded dissimilarities to the target by reassigning *N* = 1–9 randomly chosen target slots to a different value such that no lure coincided with a studied target. When reassigning all *N* = 10 slots, this procedure constructs an unrelated foil probe that shares no features with its parent target.

#### Training and test

We trained 25 networks (independent random weight initializations and stimulus orders). On each study trial, a target’s features, category, and identity code were clamped and the network learned their associations. Training continued until the network reproduced at least 80% of the identity codes correctly, reached after approximately 9-10 epochs.

Each trained network was tested on 495 probes: the 45 studied targets and, for each target, nine related lures and one unrelated foil. Target–lure similarity was defined as 1 − *N/*10. On each test trial the probe’s features were clamped and the identity layer was freed. The network’s task was to pattern-complete the corresponding studied item’s identity code.

To measure target-identifying information in each region’s trained activity, we constructed a decoder for each network and region using the 45 target settles as references. Each lure was assigned to the target whose reference pattern had the highest cosine similarity to its settled activity. Decodability was the proportion of lures assigned to their parent target.

#### Recall scoring and rejection routes to memory discrimination

The recalled identity was the CodeName column with the greatest summed activation. Following prior per-unit completion criterion *(13)*, a recall was counted only if at least half of the recalled identity’s six code units were active above a completion threshold of 0.5. This threshold corresponds to the valley of the bimodal unit-activation distribution (inactive units near 0, completed units near 0.7–0.9). Completions failing this criterion were scored as retrieval failures (recall score of 0). For counted recalls, the recall-to-reject score was (match − mismatch)*/*10 ∈ [−1, 1], comparing the recalled item’s canonical features with those of the probe. A probe was judged “old” if its recall score exceeded the network’s decision criterion (the midpoint between its mean target and mean lure recall scores) and rejected as “new” otherwise. Because retrieval failures score 0, below every network’s criterion, correct rejections can arise from both mismatching recall (recall-to-reject) and retrieval failure.

Within each network, the hit rate *H* was computed over the 45 targets and a false-alarm rate *F*_*s*_ over the 45 lures at each similarity level *s*. Discriminability was *d*^*′*^(*s*) = *z*(*H*) − *z*(*F*_*s*_). *d*^*′*^ was computed per network and averaged (mean ± SEM across the 25 networks). Pattern separation was assessed as in the original framework by comparing, for each lure, the cosine similarity between the lure’s and its parent’s settled activity in each region against their input feature overlap.

#### Cortex-only simulation

To isolate the cortical compression mechanism from the hippocampal circuitry, we simulated a single cortical layer, where one input layer projects fully to one hidden layer with kWTA inhibition. This simulation learned by the pure Hebbian dynamics, such that any compression reflects passive statistical learning of the input correlations. These dynamics lead the layer’s weights spanning the principal components of the inputs *(71)*. The network was trained on synthetic stimulus sets whose redundancy we controlled directly. Inputs were generated as combinations of a predetermined set of latent dimensions, manipulating the dimensionality of the input. We tested learning dynamics under different levels of hidden layer inhibition (*G*_*i*_). Compression was quantified as the ratio of input to hidden-layer participation ratio.

### Signatures of lossy compression

#### Lossiness from a simulation of memory retrieval and identification

To compute lossiness for all models, except for the *β*-VAE, we adapted a previously published information theoretic algorithm *(43)*. The algorithm infers a loss function based on efficiently coding (compressing) an input *X*_*target*_ into an output *X*_*lure*_. In that work, the lossiness of compression explained when two perceptually similar items are confused (generalized) or discriminated across many sensory modalities, such as a range of tones, colors, line lengths, and shapes.

In the context of the mnemonic similarity task, we determine the perceptual loss when compressing *X*_*targets*_ and *X*_*lures*_ in latent space. To simulate the cost of remembering *X*_*targets*_ given *X*_*lures*_, we linearly interpolate between *X*_*target*_ and *X*_*lure*_ in latent space. Similar images are thought to exist on a low-dimensional manifold in this latent space *(93, 96)*, and linearly interpolating between points on this manifold can be thought of as a trajectory along the manifold’s surface from one image to another. This trajectory represents samples that transform according to a primary factor of variation, such as the rotation leftward versus rightward a face, the shape of a square to circle, and the color of a blue to orange object.

After generating intermediate representations along this manifold, we calculate the cosine distance between all pairs of latent representations. Then, we convert this distance matrix to a similarity matrix and normalize values by the the maximum similarity across latent representations. Finally we simulate a confusion matrix that represents the perceptual information channel, where rows are input representations and columns are output representations, based by converting all values of the similarity matrix to probablities marginalizing over the output columns. Drawing 250 samples from this probability matrix simulates how often an input representation along the target-lure image manifold is mistaken for each output representation. Previously published R code was used to fit a model based on rate-distortion theory to the confusion matrix.

Specifically, the lossiness was inferred using the Blahut algorithm for computing an optimal, capacity-limited information channel *(163)*. This optimization procedure is grounded in rate-distortion theory and iteratively adjusts conditional probabilities to minimize the lossiness at a given information rate. Model fitting was performed over 100 iterations, ensuring convergence to a stable solution, by maximizing the log-likelihood of the observed confusion matrix under the inferred probability distributions, incorporating a prior over the lossiness that penalized deviations from symmetry and constrained diagonal elements. To improve robustness, multiple fitting attempts were conducted, selecting the best fit based on the highest log-likelihood value. This procedure returns a cost matrix where each element is the lossiness or cost of error for confusing an input (row) with each output (column element). The lossiness used in the current paper is the average lossiness across the cost matrix per pair of target and lure images.

#### Optimal lossiness from variational methods

In a *β*-VAE, the loss function is defined with a trade-off that parallels rate-distortion trade-offs. Recall that the objective function in Equation 4 is *L*_*β*-VAE_ = reconstruction error + *β* KL(*q*_*ϕ*_(*z*|*x*) ∥ *p*(*z*)). Rate-distortion functions can be approximated by interpreting the reconstruction error (here defined as the binary cross entropy loss) as the distortion *D*, the information-limiting KL-divergence term as the rate *R*, and *β* as a Lagrange multiplier that scales the regularization. Larger *β* enforces more constraints on the information channel.

This objective is a special unsupervised case of the supervised information bottleneck method which separates task-relevant and irrelevant information *(62)*. Formulating the objective function in this way enables a controllable rate-distortion trade-off:

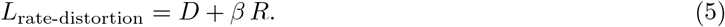

This rate-distortion loss function characterizes how a better lossy compression minimizes both the distortion and the information rate needed to transmit the source. Here, increasing *β* enforces more conservative information rates, or more aggressive compression characterizing a more capacity-limited information channel. Using this method, the lossiness of each image was measured as the average reconstruction error of the target and lure images across *β* for the seed with the lowest train/test loss. The normalized rate was measured by calculating the slope from a linear function that best fit the rate-distortion function plotted in a semi-log plot: 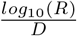.

#### Dimensionality

The first signature of lossy compression we investigated treats dimensionality reduction as a form of lossy compression, because reconstructing an input using a reduced number of dimensions introduces distortions. The base quantity is representational dimensionality, calculated as the participation ratio (PR) of the co-variance of a region’s settled activity patterns across probe trials,

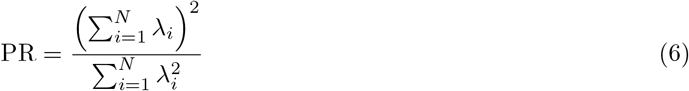

where *λ*_*i*_ are the eigenvalues of the covariance matrix computed over probes and *N* is the number of units in the region. Participation ratios are like a continuous and normalized version of the discrete principal components that explain all variance in the data. Higher participation ratios indicate neural representations with greater dimensionality and a larger capacity to represent more features or stimuli *(118, 164, 165)* which may support the separability of distinct inputs or input categories *(5, 96)*. In contrast, a lossy compression account of pattern separation predicts that separability is supported by lower dimensionality, selectively discarding less relevant features to only retain essential information.

The participation ratio is computed from a representation’s own eigenvalue spectrum, so a representation that discards most of its variance (most of its principal components) can nevertheless be measured as having a high dimensionality if the remaining variance is evenly spread. Because lossy compression is precisely a claim about variance being discarded, we anchored the measure to the network’s input variance. For example, in analyzing the C-HORSE model’s activity, the anchored participation ratio (APR) replaces the denominator with the eigenvalue spectrum of the clamped entorhinal feature layers over the input probe,

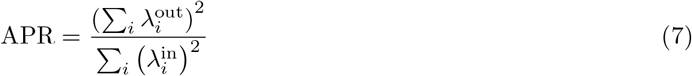

where *λ*^out^ are the eigenvalues of the region’s activity covariance and *λ*^in^ those of the input covariance.

The signature of lossy compression is a dimensionality reduction for correctly rejected versus falsely recognized lures. Because the two groups differ in size and composition of item similarity, the correct-rejection group was repeatedly subsampled to match the false-alarm group’s per-*N* similarity and the total sample size with 100 resamples.

#### Information capacity

The second signature of lossy compression we investigated is the information capacity of the representations: the number of bits a region’s activity distribution could carry under Gaussian information channels with a fixed noise level *(73)*,

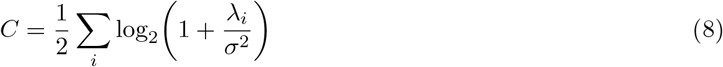

where *λ*_*i*_ are the eigenvalues of the region’s activity covariance and *σ*^2^ is the noise variance. For analysis of the C-HORSE model’s activity, because the networks settle deterministically, *σ*^2^ is selected such that the input’s capacity equaled the input’s target content, log_2_(45 total targets) bits. Capacity is thus expressed relative to an input channel with optimal efficiency for the task. The same *σ*^2^ was applied to all regions and both outcome groups within a network, so group differences reflect the representations’ eigenvalue spectra rather than the noise convention.

Unlike decodable stimulus information, capacity is not bounded by the input such that the sparse expansion recoding can exceed the input’s capacity. This does not violate the data processing inequality as it does not create stimulus information but instead indicates that more information can be represented. A lossy compression account predicts lower capacity for correctly rejected than falsely recognized lures, computed with the same procedure as the dimensionality analysis.

### MRI data acquisition

#### Kirwan dataset

This dataset included neuroimaging performed on a Siemens 3 Tesla TIM Trio scanner using a 32-channel receive-only head coil. Structural images were acquired using a T1-weighted MP-RAGE sequence with the following parameters: 192 interleaved slices; total acquisition = 8:55 min; TR = 20 ms; TE = 4.92 ms; flip angle = 25°; field of view = 256 mm; slice thickness = 1 mm; voxel resolution = 1.0 × 1.0 × 1.0 mm; 1 average. A T2-weighted pulse sequence was acquired with the following parameters: 35 interleaved slices; total acquisition = 9:44 min; TR = 6,000 ms; TE = 64 ms; flip angle = 129°; field of view = 200 mm; slice thickness = 2 mm; voxel resolution = 0.4 × 0.4 × 2.0 mm; averages = 2. This scan was positioned perpendicular to the longitudinal axis of the hippocampus for each participant prior to acquisition.

High-resolution functional images were acquired using gradient-echo echoplanar, T2*-weighted pulse sequence utilizing a multiband (MB) technique *(166)*; six functional scans were conducted that coincided with six blocks of the task. These scans had the following parameters: 72 interleaved slices; TA = 6:25 min; TR = 875 ms; TE = 43.6 ms; flip angle 55°; field of view = 180 mm; slice thickness = 1.8 mm; voxel resolution = 1.8 × 1.8 × 1.8 mm; MB factor = 8; measurements = 428. These scans were aligned with the longitudinal axis of the hippocampus for each participant. The first four TRs of each run were discarded to allow for T1 equilibration.

#### Bennett dataset

Imaging data were collected at the Center for Advanced Neuroimaging at the University of California, Riverside, on a 3 Tesla Siemens Prisma scanner with a 32-channel receive-only head coil *(38)*. Structural images were acquired using a T1-weighted MP-RAGE sequence (TR/TE = 2,400/2.72 ms; field of view = 256 × 256 × 208 mm; flip angle = 8°; voxel resolution = 0.8 × 0.8 × 0.8 mm). Functional images were acquired in two runs (study and test) using an echo-planar imaging sequence (TR/TE = 1,750/32 ms; field of view = 221 × 190.4 mm; flip angle = 75°; voxel resolution = 1.7 × 1.7 × 1.7 mm; 72 slices with no gap; anterior–posterior phase encoding; GRAPPA factor = 2; MB factor = 3).

All stimuli were presented for 2.5 s with a 0.5 s inter-stimulus interval, and left-/right-facing arrow trials requiring direction judgments were randomly interspersed as a non-mnemonic baseline. Unlike the three-choice (old/similar/new) format of the Kirwan dataset, correct rejection of a lure here corresponds to a “new” response and a lure false alarm to an “old” response.

### fMRI preprocessing

#### Kirwan dataset

MRI data were analyzed using the Analysis of Functional NeuroImages (AFNI) software (version 22.1.09; *(167)*) following the first steps of a prior publication using the same data *(37)*. Given the short acquisition time, functional scans were not slice-time corrected. Motion correction for functional scans was calculated to align each volume to the single volume of the experiment with the smallest number of outlier voxel values. Structural scans were also aligned with the minimum-outlier functional volume. Rotated structural scans were then skull-stripped and warped into MNI space using a nonlinear diffeomorphic transformation. The motion correction and MNI normalization spatial transformations were concatenated and applied to the functional scans in a single step, thus resulting in a single spatial transformation for functional data. Functional data were scaled by the mean of the overall signal for each run. No blurring was done to functional data as part of preprocessing and spatial resolution of functional scans was maintained at 1.8 mm^3^.

Using this minimally pre-processed data, we performed trial-wise GLM estimation using the GLMsingle toolbox implemented in Python *(168)*. This toolbox identifies an optimal HRF from a library of 20 canonical functional forms, performs automated denoising using data-driven nuisance regressors from selected repeat trials, and implements fractional-ridge regularization to regularize estimates based on voxel-wise reliabilities *(169)*. This pipeline does not benefit from regressing out putative nuisance components of fMRI data, such as motion, white matter, or cerebrospinal fluid, prior to running GLMsingle because those steps can bias the data-driven learning of nuisance components which potentially overlap with signal components of interest. For these reasons, our pre-processing departed from the steps in the prior work using this dataset *(37)*, i.e., we also did not use the motion scrubbing (framewise censoring of TRs with greater than 0.3° of rotation or 0.6 mm of translation in any direction, as well as the immediately preceding TR) or coverage masks removing very low EPI signal.

Region of interest maps were created for the ventral visual stream (IT), as well as the hippocampal subregions (DG/CA3) that classically supports pattern separation for inputs from the object processing stream. The IT mask was obtained from the Harvard-Oxford atlas *(170)* and the DG/CA3 mask was the same as in prior work *(37, 171)*. Somatosensory masks were used as experimental controls and were obtained from the Juelich histological atlas *(172)*.

For each subject, we masked all ROIs. Then, for 1 of the 6 runs, we split the 2nd run for each participant into two half runs because GLMsingle requires repeat trials across runs for cross-validation and the original task only includes repeat trials within a run. With stimulus duration = 2.5 s and repetition time = 0.875 s, we ran GLMsingle with default options. Outputs used for further analysis included the per-trial evoked activity maps and map of HRF indices. Split-half reliability on *β*s were computed voxelwise.

#### Bennett dataset

MRI data were preprocessed with fMRIPrep 25.2.5 *(173)* using default settings. Anatomical preprocessing included intensity non-uniformity correction, skull-stripping, and nonlinear spatial normalization to the MNI152NLin2009cAsym template. Functional preprocessing included head-motion estimation and correction, susceptibility distortion correction using the opposite phase-encoded image pairs, boundary-based co-registration to the T1-weighted image, and resampling in a single interpolation step to template space at 2 mm isotropic resolution. As with the Kirwan dataset and recommended for the downstream GLMsingle pipeline *(168)*, no spatial smoothing was applied and no nuisance time series were regressed so that noise dimensions could be determined in a data-driven manner. Prior to single-trial modeling, each run’s preprocessed BOLD time series was converted to percent signal change on a voxel-wise basis, expressing each time point as the percent deviation from that voxel’s within-run temporal mean.

Trial-wise GLM estimation again used the GLMsingle toolbox *(168, 169)*, fitting the study and test runs jointly with stimulus duration = 2.5 s and repetition time = 1.75 s and default options. Each of the 256 unique object stimuli was modeled as a separate condition with an onset-locked regressor; the 64 target stimuli were modeled as the same condition at their study and test presentations. In contrast to the Kirwan dataset, these repeated targets naturally furnished cross-run repeat trials for GLMsingle’s cross-validation, so no run splitting was required. Outputs used for further analysis included the per-trial evoked activity maps, the map of HRF indices, and voxel-wise cross-validated variance explained (*R*^2^).

The same set of regions of interest was defined for the Bennett dataset. Medial temporal lobe subregions (anterolateral entorhinal cortex, DG/CA3, CA1, and subiculum) were segmented per participant using DeepFLASH *(174)* applied to the fMRIPrep-preprocessed T1-weighted image in native space, warped to template space (2 mm) using the fMRIPrep-derived anatomical transform with label-preserving interpolation, and combined across hemispheres into bilateral masks. The IT cortex as well as sensorimotor control regions were defined at the group level in template space with the same procedure as the Kirwan dataset. For all analyses of both datasets, voxels within each region of interest were retained if their GLMsingle cross-validated *R*^2^ exceeded 10%.

### Representational similarity analysis

#### Obtaining diagnostic stimulus features to compress

For each lure pair (target image *X*_*target*_ and lure image *X*_*lure*_), we extracted convolutional neural network representations from AlexNet’s layers as this was the most predictive model behaviorally. We train using all of AlexNet’s layers but only analyze conv5, the deep layer most analogous to higher-order ventral visual cortex IT *(59, 60)*. To identify the information most diagnostic of each image’s object, channel activations were weighted by Grad-CAM (Gradient-weighted Class Activation Mapping). This is an explainable AI technique that highlights what information within each image was most important to recognize the object within the image.

For each image, the logit of its top-1 predicted class (e.g., sports car, golden retriever, cheeseburger) was backpropagated to each layer, channels were weighted by their positive gradient values, and the weakest 50% of channels were set to zero. To suppress artifactual responses arising from zero-padding at the image border, the outermost pixels of each feature map were cropped prior to spatial pooling. Spatially pooled z-scored feature vectors served as model inputs.

#### Compression model implementing redundancy reduction and information bottlenecks

We trained a single encoder-decoder network separately for each fMRI dataset on that dataset’s stimulus set (85%/15% train/test split). The training objective jointly combines a reconstruction term, an information bottleneck, and a Barlow-Attneave redundancy-reduction constraint:

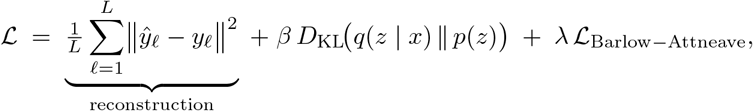

where *L* is the number of AlexNet layers (*L* = 6; conv1–conv5 and fc7), *y*_*ℓ*_ is the reconstruction target for layer *ℓ* (the absolute channel-wise difference between the pair members’ Grad-CAM–weighted feature vectors), *ŷ*_*ℓ*_ is the decoder’s reconstruction of *y*_*ℓ*_, *x* denotes an image’s input features, *z* is its latent code, *q*(*z* | *x*) is the encoder’s posterior over the latent code, *p*(*z*) = *N* (0, *I*) is the standard Gaussian prior, *D*_KL_ is the Kullback–Leibler divergence. The *β* is the information bottleneck weight that determines the strength of information compression and *λ* weights the Barlow-Attneave redundancy-reduction term L_Barlow_.

The encoder mapped each image’s Grad-CAM–weighted features to a latent code *z* (32 dimensions per layer) parameterized as a diagonal Gaussian *q*(*z* | *x*) = *N* (*µ, σ*^2^). For each AlexNet layer, the encoder comprised a 64-unit linear projection (no bias), a 64-unit ReLU hidden layer, and a linear head producing the 32-dimensional mean and log-variance of the latent code (log-variance bias initialized at −2 and clipped to [−10, 10]). The decoder received the concatenated per-layer latent contrasts 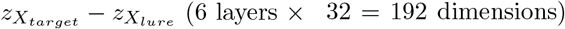, passed them through a shared 64-unit ReLU layer, and reconstructed the absolute channel-wise feature difference 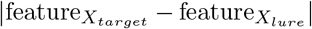 between pair members through per-layer linear output heads matching each layer’s feature dimensionality, so that reconstruction targeted the features that distinguish a target from its lure. The network was trained using the Adam optimizer with a learning rate of 10^*−*4^ and gradient-norm clipping at 1.

The redundancy-reduction term implements the Barlow-Attneave efficient coding *(42, 65, 101)*. Two distorted views of the same stimulus (random resized crop, color jitter, horizontal flip, Gaussian blur, applied at the pixel level) were passed through the frozen AlexNet feature extractor and the shared encoder. Redundancy reduction is imposed by penalizing the cross-correlation matrix *C* of the batch-standardized *B* latent codes toward the identity matrix:

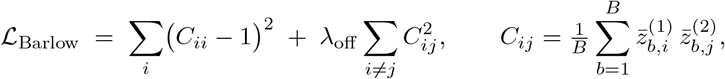

where on-diagonal *C*_*ii*_ terms enforce invariance of each latent dimension (analagous to putative IT cortex function *(93)*) to the distortions and off-diagonal *C*_*ij*_ terms minimize redundancy across dimensions *z* (analagous to Barlow-Attneave redundancy reduction *(42, 101)*). The chosen *λ* = 0.005 was calibrated to approximately equate the redundancy-reduction term’s loss with the reconstruction term’s loss at initialization.

To implement the information bottleneck *(66)*, during training, *β* was held at 10^*−*5^ for 300 pretraining epochs and then increased log-uniformly over 3,000 epochs with model snapshots saved at each *β*. This yields a family of compressed codes spanning effectively uncompressed to aggressively and overly compressed representations. The range of *β* values considered were determined by assessing if the mean Bhattacharyya distance had collapsed below 1% of its uncompressed value (*d*^*′*^ ≈ 0.8), suggesting degenerate codes. The analyzed range comprised 7 compression levels spanning uncompressed (*β* = 1.5 × 10^*−*4^) to strongly but non-degenerately compressed codes (*β* = 3.7 × 10^*−*2^ with a *d*^*′*^ ≈ 2).

#### Model representational dissimilarity matrix

Because each stimulus code is a multivariate Gaussian, the discriminability of a target-lure pair was quantified as the Bhattacharyya distance *D*_*B*_ between *q*(*z* | *x*_*a*_) and *q*(*z* | *x*_*b*_), a generalized analog of *d*^*′*^ in signal detection theory. Pairwise distances were averaged within each stimulus similarity bin (six bins in the Kirwan dataset; five in the Bennett dataset) to obtain a model discriminability profile across bins at each compression level.

#### Neural representational dissimilarity matrix

For each subject, ROI, and stimulus similarity bin, neural discriminability was computed as the mean cosine distance between the trial-wise activity patterns (single-trial activity estimates) evoked by each target and its corresponding lure, averaged over pairs within the stimulus similarity bin. Analyses were restricted to trials with correct lure responses (“Similar” in the Kirwan dataset *(37)* three-alternative task and “New” in the Bennett dataset *(38)* two-alternative task). Subjects lacking correct trials in any bin were excluded, yielding *n* = 37 (Kirwan dataset) and *n* = 54 (Bennett dataset). A benchmark analysis included all trials regardless of accuracy (*n* = 48 and *n* = 61).

#### Representational similarity in regions of interest

For both model and neural representations, we compared the representational dissimilarity matrices over similarity bins using Mantel tests. This calculates the Pearson correlation with a permutation of bin labels at the subject level because distance data is non-independent. Group-level significance at each compression level was assessed by one-sample *t*-tests on subjects’ Mantel *r*-values against zero and *p*-values were adjusted for multiple comparisons using FDR (Benjamini-Hochberg) correction over ROIs (IT and DG/CA3) and compression levels.

Primary motor and somatosensory cortex were considered as experimental control regions to provide a baseline. In exploratory analysis (see **Supplementary Results**), we additionally analyze a subregion of the anterior temporal lobe, alEC, which was previously implicated in memory discrimination *(171)*. alEC is also of interest because it is part of the anterior temporal lobe, a key semantic hub that may be associated with the process of semanticization: transforming memory representations into a more abstract, gist-like representations that loses sensory details *(88, 175)*. We compare the individual alEC parcels to a recent parcellation of the anterior temporal lobe and find that there is a complete spatial overlap between the regions’ voxels *(176)*.

### Statistical analysis

To test the effect of lossiness on the ease of memory discrimination, we used a Spearman correlation between lure bin and lossiness. To test the effect of lossiness on memory discrimination, we tested a linear model with mixed effects at the trial-level nested by participant:

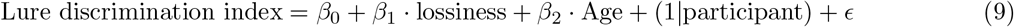

The lure discrimination index was calculated across trials within each lure bin. The lossiness was the mean across the images presented in each lure bin. We only included age as a covariate for the dataset that spanned the lifespan and the dataset containing children and adolescents. Minimally accounting for the linear effects of general age-related phenomena is important because there are numerous differences related to ongoing neurodevelopment with heterogeneous effects across regions. The other key purpose of this covariate was to control for known aging effects on the Mnemonic Discrimination Task, where performance is substantially lower in older ages *(26)*. Finally, the effects of age are often non-linear. Neurodevelopmental processes related to functional connectivity (statistically related to the covariance matrix we use to calculate neural dimensionality) develop non-linearly and the shape of the non-linearity depends on the particular brain region *(177)*. To test both linear and non-linear effects of age on the relationship between lossiness and memory discrimination, we used generalized additive models with penalized splines, a method which allows for statistically rigorous modeling of linear and nonlinear effects while minimizing over-fitting *(89)*. We tested the model:

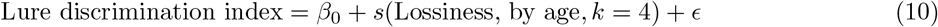

The model included *k* as a smooth term for lossiness, capturing non-linear effects of lossiness across different age groups, using 4 basis functions and fit using restricted maximum likelihood estimation and fixed effects.

### Ethics statement

All procedures were approved by the institutional review board of the university at which the data were collected: online data collection under IRB #20216528 at the University of California - Irvine; child sample under IRB #2021-5654 at New York University; undergraduate sample under IRB # 2008-6128 at the University of California - Irvine; the undergraduate and fMRI sample under IRB X14208 at Brigham Young University; and the fMRI sample at the University of California - Riverside under IRB # 2016-097.

### Data, code, and materials availability

All data and code needed to evaluate and reproduce the results in the paper are present in the paper and/or the Supplementary Materials. This study did not generate new materials and drew on previously published data. The code used for modeling and analysis will be deposited to Zenodo upon publication. The Kirwan fMRI dataset is available at OpenNeuro.

### Citation diversity statement

Recent work in several fields of science has identified a bias in citation practices such that papers from women and other minority scholars are under-cited relative to the number of such papers in the field *(178–186)*. Here we sought to proactively consider choosing references that reflect the diversity of the field in thought, form of contribution, gender, race, ethnicity, and other factors. First, we obtained the predicted gender of the first and last author of each reference by using databases that store the probability of a first name being carried by a woman *(182, 187)*. By this measure (and excluding self-citations to the first and last authors of our current paper), our references contain 12.27% woman(first)/woman(last), 12.63% man/woman, 21.27% woman/man, and 53.84% man/man. This method is limited in that a) names, pronouns, and social media profiles used to construct the databases may not, in every case, be indicative of gender identity and b) it cannot account for intersex, non-binary, or transgender people. Second, we obtained predicted racial/ethnic category of the first and last author of each reference by databases that store the probability of a first and last name being carried by an author of color *(188, 189)*. By this measure (and excluding self-citations), our references contain 6.18% author of color (first)/author of color(last), 12.90% white author/author of color, 22.74% author of color/white author, and 58.18% white author/white author. This method is limited in that a) names and Florida Voter Data to make the predictions may not be indicative of racial/ethnic identity, and b) it cannot account for Indigenous and mixed-race authors, or those who may face differential biases due to the ambiguous racialization or ethnicization of their names. We look forward to future work that could help us to better understand how to support equitable practices in science.

## 4 Acknowledgments

D.Z. acknowledges funding from the George E. Hewitt Foundation for Medical Research. A.M.B. acknowledges funding from the National Institute on Aging (R21AG072673 and R01AG088306). M.Y. and A.M.B. acknowledge funding from the National Institute for Mental Health (R01MH128306). We thank Ilana J. Bennett for the fMRI dataset including older adults (collected under K99AG047334). We thank Dhairyya Singh and Kenneth A. Norman for helpful discussions and feedback, and anonymous reviewers for their constructive comments.

