## Supplementary Materials for "A compressed code for memory discrimination"

October 15, 2026

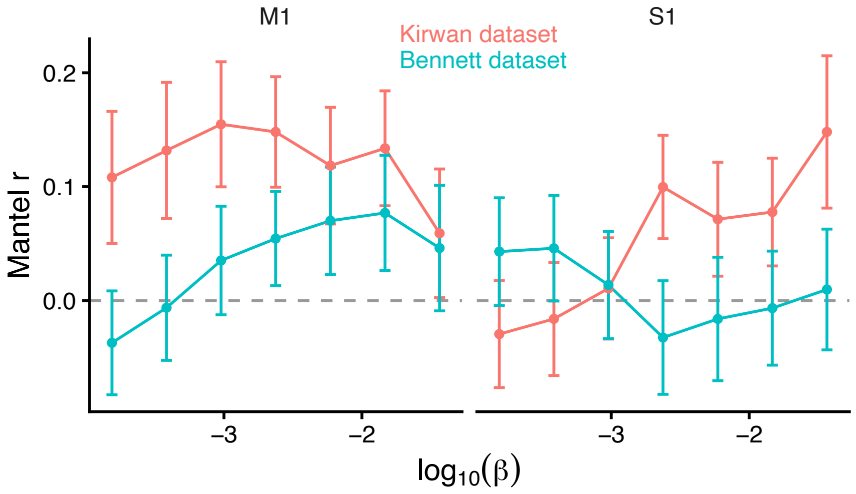


Figure S1: Null results for motor and somatomotor regions M1 and S1 in the representational similarity analysis. $\beta$ is the strength of compression. No effect survives FDR-correction.

### Supplementary Results

#### Representational similarity analysis

Control regions showed no significant RSA correspondence in either dataset. We evaluated whether compression was confounded by a global or task-general signal by analyzing motor and somatomotor regions putatively uninvolved in the compression or expansion process (**Figure S1**. In the Bennett dataset, where IT reached significance, M1 and S1 were uniformly null (all $p_{\mathrm{FDR}} > 0.05$), indicating that the IT correspondence does not reflect a global or task-general signal. In the Kirwan dataset, M1 and S1 did not reach significance but showed a few trending effects (minimum $p_{\mathrm{FDR}} = .06 - .07$) with the strongest correlation ($r = 0.16$) at the highest level of compression being numerically comparable to the hypothesized-regions' correlation magnitudes. It is feasible that some motor-related or preparatory motor compression is implicated in making the lure discrimination decisions. However, this trending relationship did not replicate in the other dataset.

An exploratory analysis of alEC showed nominal positive correspondences in both datasets (**S2**), though none survive multiple comparisons within this single region across levels of compression. The minimum $p_{\mathrm{FDR}} = .10$ and $.19$ for the Kirwan and Benentt datasets, respectively.

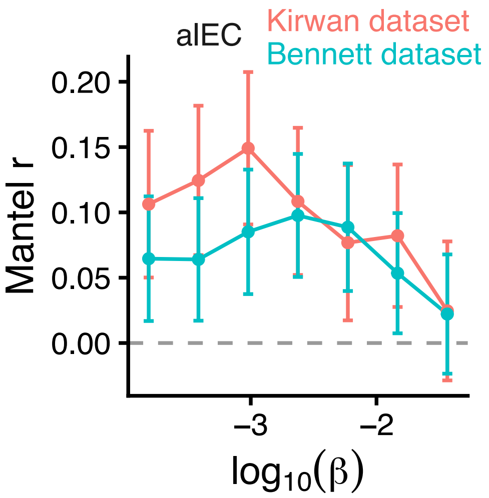


Figure S2: Null results for aIEC in the representational similarity analysis. $\beta$ is the strength of compression. No effect survives FDR-correction.

#### Predicting lure false alarms

Individuals tended to have worse pattern separation performance on images with more aggressive compression. Consistent with the compression hypothesis, greater lossiness correlated with lower lure false alarms (**S3**). This relationship largely replicated in 20 out of 24 tests across 4 datasets and the perceptual and semantic features tested. However, the effect of the normalized rate was less consistent, suggesting that lossiness is a more robust metric.

#### Neural network learning curves and generalization test

Training curves for the autoencoder show that the divergence between training and validation loss is modest in absolute terms and, importantly, the validation loss converges (**S4**), indicating that the model is not progressively fitting noise during training at the expense of held-out performance. We trained an autoencoder with a relatively small data set of experimental stimuli, which may be prone to overfitting on those images. This may confound compression results that depend on the image encodings. We therefore aim to compare our autoencoder to an autoencoder that was trained in a broadly similar way as the other models we considered (e.g., VGG-16) which used a large dataset of naturalistic images that were not also used in the human experiments. To assess whether our task-trained autoencoder captured representational structure comparable to a network trained on naturalistic image statistics, we compared our autoencoder's 256-dimensional image embeddings to those of a VGG-16 autoencoder pretrained on ImageNet [1]. The lat-

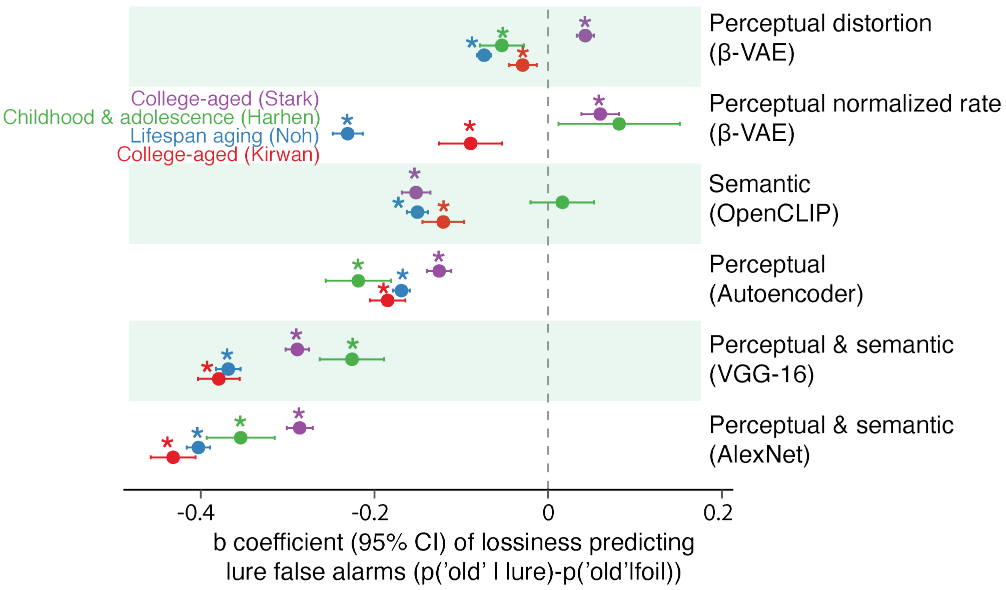


Figure S3: Consistent with our hypothesis, greater lossiness across people is related to reduced lure false alarms. This suggests that redundancy reduction can decrease overlap across an individual's memories, but over-compression can produce false memory.

ter autoencoder was not trained on the stimuli used in the memory discrimination task. Using each network to encode task stimuli, we calculate each network's pairwise cosine-distance between all stimuli to generate a image $\times$ image representational dissimilarity matrix. Then, we perform representational similarity analysis between the image matrices, finding that they were strongly correlated (Spearman $\rho = 0.78$–$0.80$, Pearson $r = 0.77$–$0.80$, all $p < .001$), indicating that the two networks organized the stimuli in highly similar ways in their latent spaces despite differences in architecture, training diets, and latent dimensionality. Latent-space interpolations between each target–lure pair were likewise nearly identical across networks, with the step-wise distance matrices correlating at $\rho = 0.99$, suggesting that the geometry of the target-to-lure recall trajectories is preserved across the two representational spaces. This geometry of the atent space is interpretable albeit lossy. Linear interpolations between a target and its lure traverse smooth intermediate reconstructions, consistent with meaningful representations learned under lossy compression. We visualize the linear interpolations that we use to operationalize memory recall from an image to its associated lure **Figure S5**.

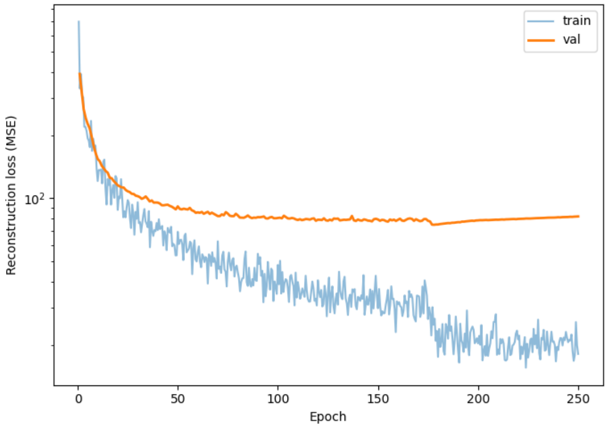


Figure S4: **Learning curves.** Reconstruction loss (squared error summed over all pixels and channels of each image and averaged across the batch) across training epochs for the autoencoder, with training loss (blue) shown per step and validation loss (orange) per epoch. Validation loss plateaus at around epoch 75 and remains stable through 250 epochs.

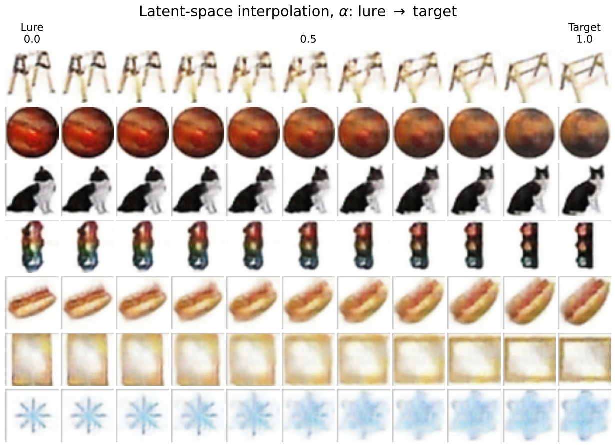


Figure S5: **Recall trajectories from model-generated interpolation** Simulated recall trajectory from a presented lure to the studied target, based on linear interpolation between learned vector representations of the lure and target stimuli, where the numeric values are $\lambda$: $z(\lambda) = z_{\text{target}} + \lambda\,(z_{\text{lure}} - z_{\text{target}})$ .

#### Simulation with MNIST

To demonstrate the intuition behind how distortion can support discrimination in practice, we first apply distortion and dimensionality metrics to a popular machine learning dataset. Does pattern separating handwritten numbers in an image dataset into digits benefit from dimensionality reduction or expansion [2]? To obtain encodings, images were decorrelated into orthogonalized components using principal components analysis. After dimensionality reduction, reconstructions of the original inputs were generated by decoding from 2 to 784 (the number of pixels) principal components and define a distortion metric as the mean squared error **(Figure S6A)**. Visualizing all of the images in a 2-dimensional space highlights how distinct digits can become more separated (using a non-linear dimensionality reduction method for illustration) **(Figure S6B)**. When fewer dimensions are used to encode $X_1$, the reconstruction $X_2$ has a larger error $d(X_1, X_2)$ **(Figure S6C)**. Expansion can reconstruct all the variance of the source but the addition of principal components has diminishing returns for information about fine details.

With these representations, we can assess the quality of pattern separation under differing levels of compression or expansion. The quality of separation is measured by the distance between clusters, the quality of clustering, and the classification accuracy. The Euclidean distance across encoding dimensions represents the separability of images in each class of digits. The Euclidean distance increases with the raw dimensionality (lighter blues), seemingly supporting the dimensionality expansion hypothesis **(Figure S6D)**. However, in high-dimensional statistical learning, the dimensionality $d$ is commonly normalized by $\sqrt{d}$ to help counter the "curse of dimensionality." The curse refers to how increasing dimensionality tends to indiscriminately expand the distances between all points, making variations in distance lower, less meaningful, and less separable in practice. Indeed, after normalizing the dimensionality, Euclidean distance decreases with increasing dimensionality. Consistent with the compression hypothesis, Euclidean distance increases with distortion incurred by reducing the number of normalized dimensions.

Next, we evaluate clustering and classification of compressed versus expanded representations. Clustering quality is measured by the silhouette score, the average intra-cluster distance to the nearest-cluster distance. A score of 1 indicates good separation, 0 indicates some overlap, and -1 indicates erroneous overlap. Classification accuracy is measured by k-nearest neighbor classification using $k = 200$, where an image is classified based on the majority class among its k-closest neighbors in the space of encoding dimensions. Consistent with the compression hypothesis, the greatest silhouette scores and classification accuracies are achievable using more compact lower-dimensional representations that incur greater distortion **(Figure S6E-F)**. Taken together, these results characterize how lossy compression can support learning separable

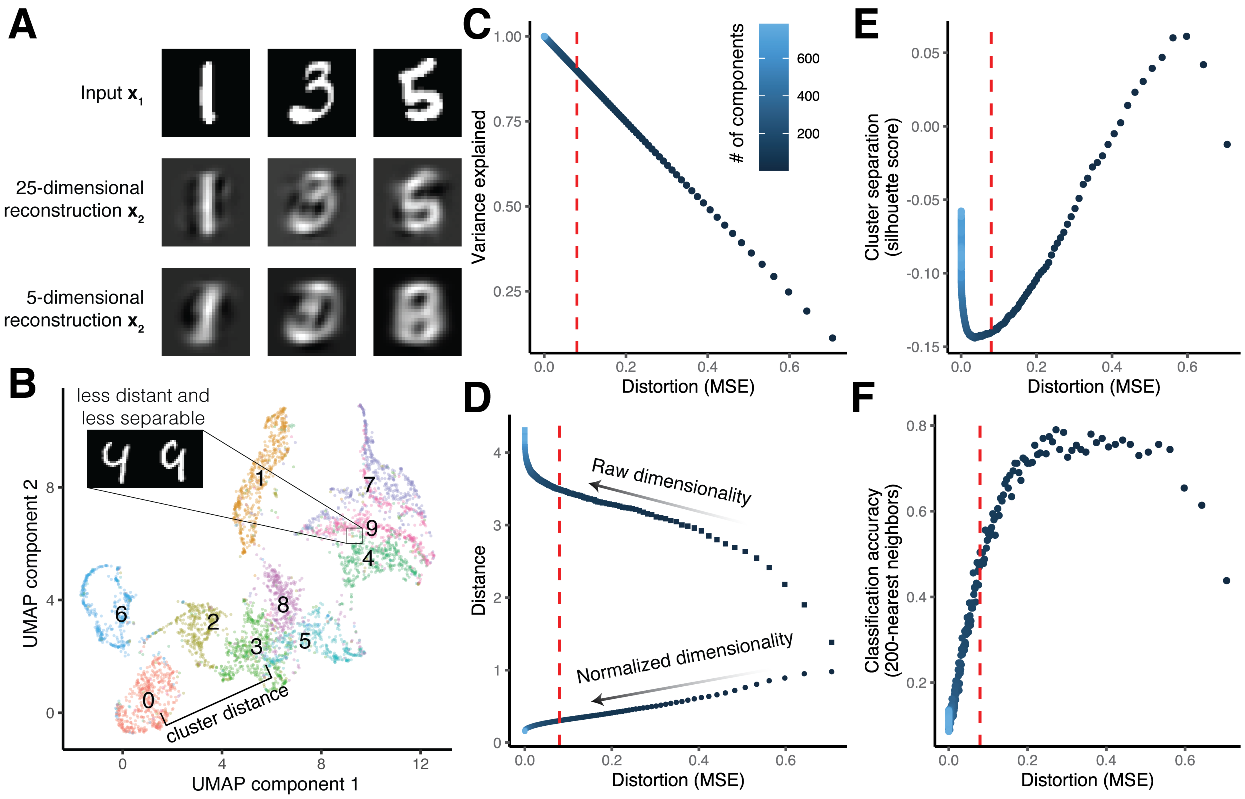


Figure S6: **(A)** Principal components analysis was used to generate a reconstruction of the images from the main 25 or 5 dimensions of variation, resulting in distortion. **(B)** A 2-dimensional embedding highlights how distant data points may be more separable and nearby data points across clusters may be more confusable. **(C)** Dimensionality reduction incurs greater distortion. The red dotted line indicates the number of components explaining over $90\%$ of variation. **(D)** The distances between clusters increase with raw dimensiona lity but decrease with normalized dimensionality. However, cluster distances increase with normalized dimensionality reduction and increased distortion, whereas distances decrease raw dimensionality reduction and increased distortion. **(E)** Cluster separation increases with distortion. **(F)** The classification accuracy increases with distortion.

representations of similar inputs, particularly when the inputs have lower intrinsic dimensionality [3].
